# Active palpation underlying shape perception is shaped by physiological thresholds and experience

**DOI:** 10.1101/2020.10.27.356584

**Authors:** Neomi Mizrachi, Guy Nelinger, Ehud Ahissar, Amos Arieli

**Author notes:** Equal contribution (Ehud Ahissar, Amos Arieli). **Corresponding authors email addresses:** Ehud Ahissar Amos Arieli Neomi Mizrachi.

## Abstract

Hand movements are essential for tactile perception of objects. However, why different individuals converge on specific movement patterns is not yet clear. Focusing on planar shape perception, we tracked the hands of 11 participants while they practiced shape recognition. Our results show that planar shape perception is mediated by contour-following movements, either tangential to the contour or spatially-oscillating perpendicular to it, and by scanning movements, crossing between distant parts of the shapes’ contour. Both strategies exhibited non-uniform coverage of the shapes’ contours. We found that choice of strategy during the first experimental session was strongly correlated with two idiosyncratic parameters: participants with lower tactile resolution tended to move faster; and faster-adapting participants tended to employ oscillatory movements more often. In addition, practicing on isolated geometric features increased the tendency to use the contour-following strategy. These results provide insights into the processes of strategy selection in tactile perception.

**SIGNIFICANCE STATMENT:** Hand movements are integral components of tactile perception. Yet, the specific motion strategies used to perceive specific objects and features, and their dependence on physiological features and on experience, are understudied. Focusing on planar shape perception and using high-speed hand tracking we show that human participants employ two basic palpation strategies: Contour-following and scanning. We further show that the strategy chosen by each participant and its kinematics depend strongly on the participant’s physiological thresholds – indicative of spatial resolution and temporal adaptation - and on their perceptual experience.

## INTRODUCTION

Perception usually co-occurs with sensors’ movements (Ahissar & Arieli, 2001; Brecht et al., 2011; Cole, 2004; Diamond et al., 2008; Halpern, 1983; Kleinfeld et al., 2006; Land, 2006; Lederman & Klatzky, 1987; Rucci et al., 2018; Schroeder et al., 2010). Moreover, in primates, it has been established that hand movements are an integral component of tactile perception of objects’ features (Ahissar & Assa, 2016; Bensmaia & Hollins, 2003; Cascio & Sathian, 2001; Gamzu & Ahissar, 2001; Gibson, 1962; Hollins & Bensmaïa, 2007; Katz, 2013; Lederman & Klatzky, 1987; Saig et al., 2012; Weber et al., 2013). Yet, the nature of these movements and their dependency on other perception-relevant factors are not sufficiently characterized. In a seminal series of studies, Lederman, Klatzky and colleagues established the initial description of active “Exploratory procedures” (EPs) of 3D objects – stereotyped movement patterns having invariant characteristics. Thus, for example, ‘Pressing’ was found to be the primary EP for evaluating hardness, ‘Lateral motion’ for evaluating texture and ‘Contour-following’ (*CF*) for evaluating the shape of 3D objects (Lederman & Klatzky, 1987). *CF* variants were further analyzed in consequent studies (Klatzky & Balakrishnan, 1991; Lederman & Balakrishnan, 1991; reviewed in Nelinger et al., 2015). Quantitative studies of planar (2D) objects revealed that humans adapt their movement patterns to the spatial characteristics of the scanned surfaces; speed, for example, is adapted to spatial frequency (Gamzu & Ahissar, 2001) and direction – to the spatial orientation of the surface’s texture (Drewing, 2012; Lezkan & Drewing, 2018). Such adaptations evolve through development (Withagen et al., 2013), can be acquired via consequent exposures to specific features (Lezkan & Drewing, 2018; Withagen et al., 2013) and can be directly taught (Gamzu & Ahissar, 2001). In these studies, the adaptations resulted in maintaining specific sensory cues in a given “working range”, likely optimal for sensation, consistent with principles of closed-loop control (Ahissar & Assa, 2016; Ahissar & Vaadia, 1990; Buckley & Toyoizumi, 2018a; Marken, 2009). Similar maintenance of “controlled variables” has been observed in other tactile tasks, both in humans and rodents (Riley et al., 2002; Saig et al., 2012; Saraf-Sinik et al., 2015; Smith et al., 2002). Overall, studies on active touch have taught us that hand movements are an integral part of tactile perception, are adapted to the touched objects and to the attended features, are learned by experience and can be taught and that they are controlled continuously during object palpation. What remains unexplained is why people differ in their scanning strategies, and how these feature-oriented strategies depend on their idiosyncratic sensory limitations and history of practice. In the present study, we address these questions in detail in a simple tactile identification task of planar (2D) shapes. We first characterized the repertoire of exploration strategies among human participants and then investigated their dependencies on idiosyncratic sensory physiology and practice history.

## RESULTS

Eleven human participants participated in this study. Hand movements were recorded in all conditions; separate finger movements were not allowed (the three palpating fingers were bound together). Nineteen objects were used, grouped in two classes (‘shapes’ and ‘features’), each including several sets (Fig. 1A). Three training protocols were used (see Materials and Methods).

**Figure 1:**
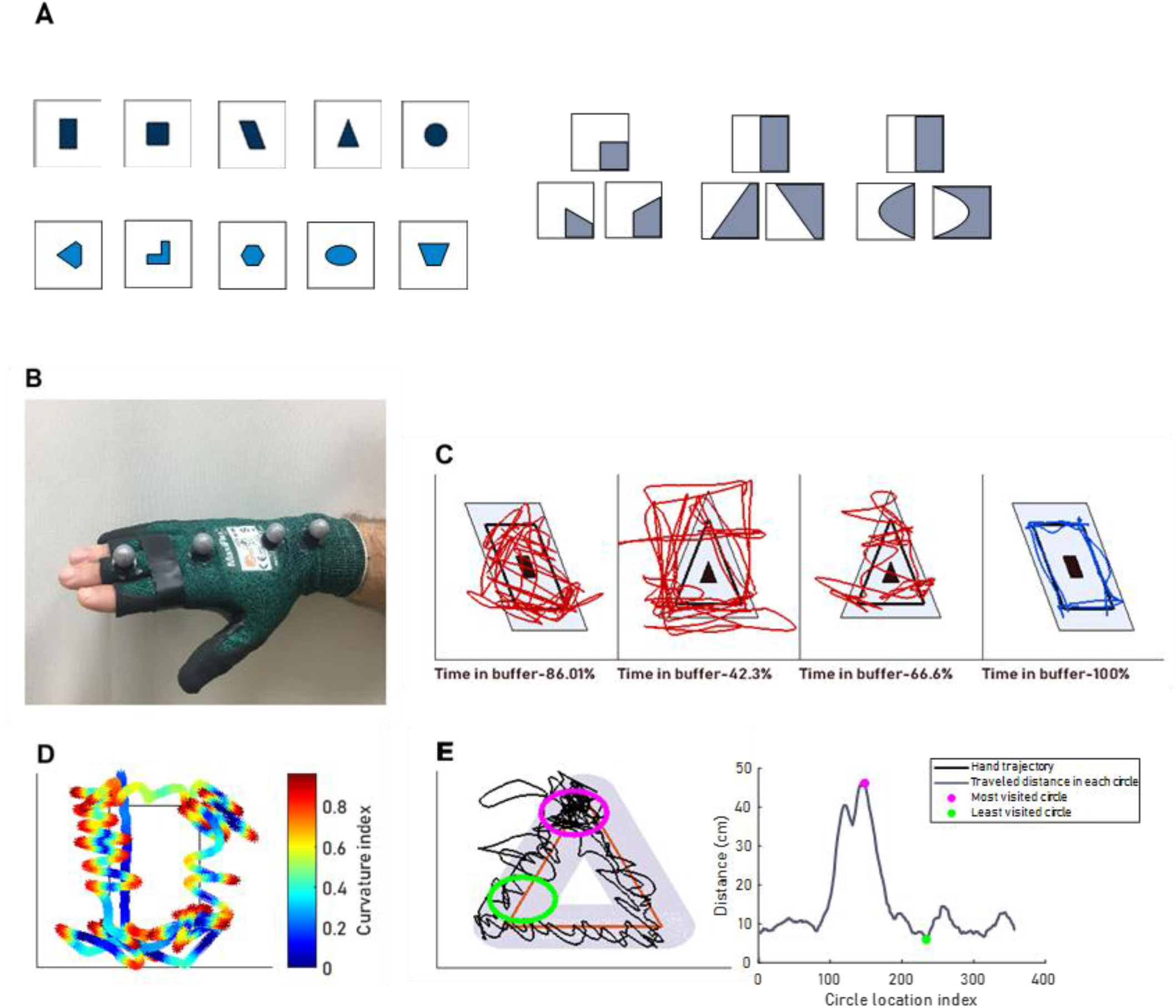
Shapes, tasks, recording and analysis. **A** - Tactile objects: Two sets of tactile objects were used Shapes (left) and Features (right). Shapes were divided into set A (top, black) and set B (bottom, blue). Features were divided into three blocks: Angle (left), Tilt (middle) and Curvature (right). For all tactile objects, the colored area was raised by 25 μm. **B** - The glove used with four designated Vicon markers attached to it. Two markers were approximately attached above the wrist carpel bones and two above middle finger’s middle and proximal phalanges. **C** - Trials classification: *Sc* trials were trials in which the hand-trajectory (red) crossed the inner object (black) or did not spend most of the time in the gray buffer, or both. *CF* trials were trials in which the hand-trajectory (blue) did not cross the inner object (black) and stayed for most of the time in the light blue buffer (left). **D** - Curvature approximation: An example trial: in each 20 mm trajectory segment, the normalized relation between shortest- and traveled-distance was calculated to evaluate the segment curvature index. **E** - An example demonstrating the calculation of a trial’s focal index. Left: overlapping circles with a radius of 10 mm (gray) were plotted on the shape’s outline (red). The traveled distance in each circle was calculated. Circles with the maximal and minimal value of traveled distance (purple and green corresponding) are marked. Right: Calculated traveled distance (y-axis) in each of the circles (x-axis). The difference between the trial’s maximal and minimal value of traveled distance (purple and green) divided by their sum is the trial’s focal index.

### Contour-following and scanning are two common procedures for planar shape recognition

In each trial, the trajectory of the hand was superimposed on the outline of the shape. Characteristic patterns were revealed and classified into two major classes: Contour-following (*CF*), where the trajectory followed the shape’s outline (Fig. 2A), and Scanning (*Sc*), where the trajectory crossed between distant parts of the shapes’ contour (Fig. 2B; see Materials and Methods). *CF* trajectories were sub-classified as *Linear* (Fig. 2A, left) and *Oscillating* (Fig. 2A, middle); trials could include one sub-class (Fig. 2A, left, middle) or both (Fig. 2A, right). *Sc* trajectories were characterized with different degrees of shapes coverage and focusing on specific shapes’ features. On average *Sc* trials differed from *CF* in their kinematics: *Sc* trials were characterized by higher tangential (scanning) speeds, longer traveled distances, higher entropy and higher speeds along the z-axis (perpendicular to the scanned surface) (Fig. 2C, p<0.005 for all differences). These exploration strategies were not specific to any of the three training protocols (Supplementary material, Fig. 2-Sp8–9). *CF* and *Sc* kinematics differences were replicated for individual protocols in most cases (Fig. 2-Sp table 1).

**Figure 2:**
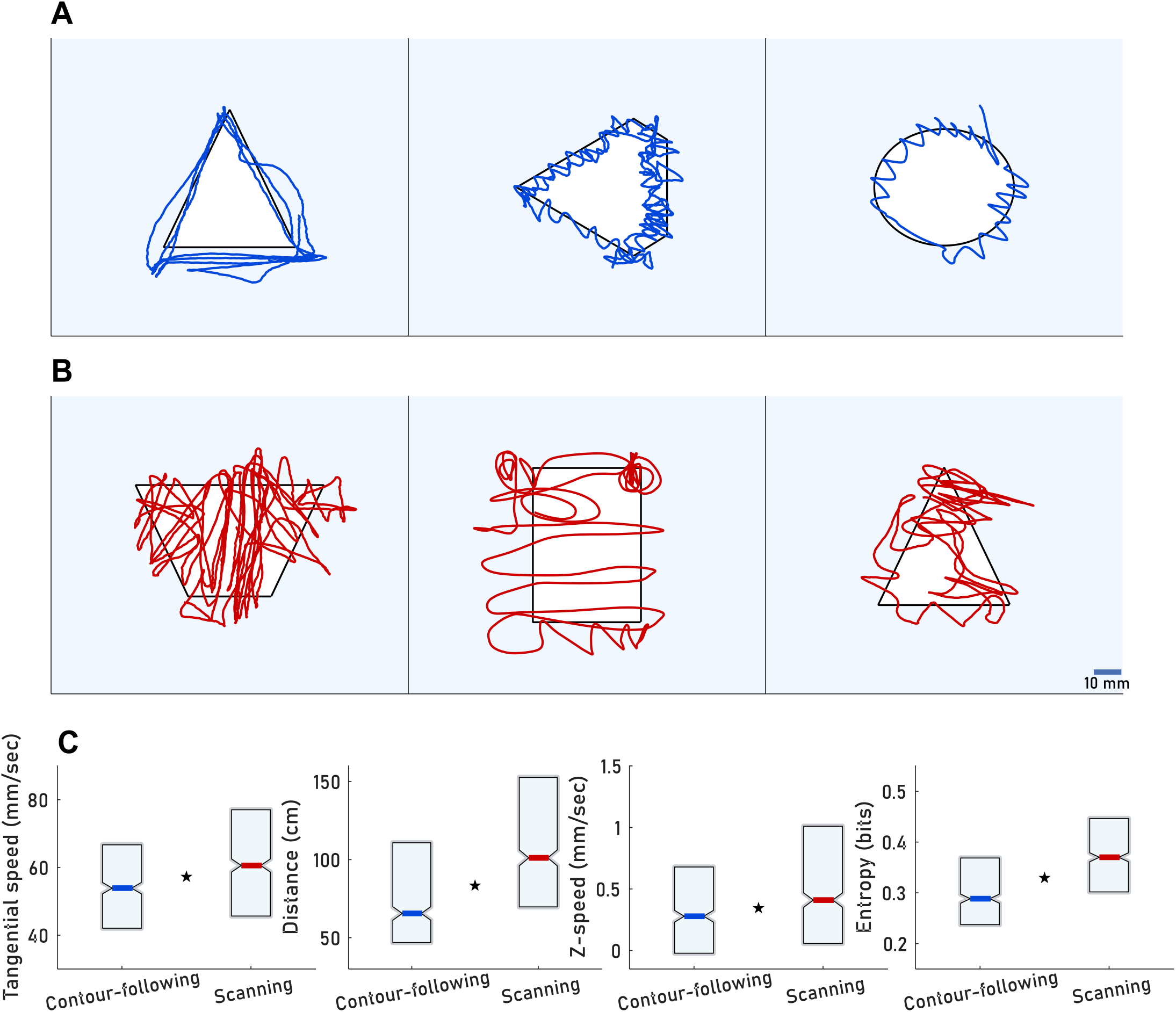
Classes of motor patterns. Strategies of hand movement were divided into subtypes and grouped to two main families: **A** - *CF* (Contour-Following) trials, which were further divided to ‘*Linear*’ (left) ‘*Oscillating*’ (middle) or both (right). **B** - *Sc* (Scanning) trials, in which the trajectory crossed between distant parts of the objects’ contour. Scale bar (B, right) corresponds to all example trials. **C** - Median and quartiles of *CF* and *Sc* kinematics (p<0.005 for all differences). N _subjects_: Eleven (training protocol I-5, II-3, III-3 subjects). N _Sessions_ =51 sessions, 4-5 sessions per participant (training protocol III - 5 sessions, I and II-4 sessions). N _trials_=1196 trials (training protocol I=417 trials, II-360 trials, III-419). N _*CF* trials_: 537 (protocol I121, II-249 III-167). N _*Sc* trials_: 659 (protocol I-296, II-111 III-252 trials). N _Trials per subject_: training protocol I: 75-90, II: 119-121, III-105-155 trials per subject.

### Focal palpation

A focal index (see Materials and Methods) was used to evaluate the degree in which the participants focused on specific parts of the explored object (Fig. 1E purple). *Sc* trials’ median focal index was significantly higher than that of the *CF* trials (Fig. 3A, p<0.005). This trend was similar between training protocols (Supplementary material, Fig. 2-Sp table 1). This difference was evident for all objects of Sets A and B (Fig. 3B). A significant correlation was observed between the median focal index and the object’s sharpest angle (excluding the circle and ellipse) for *CF* trials (Fig. 3C r = −0.86, p = 0.006, adjusted alpha level = 0.016; 0.05/3). Although not significant, a similar correlation was observed for *Sc* trials (Fig. 3D r _Pearson_= - 0.69, p_ _Pearson_= 0.057, r _Spearman_= −0.65, p_ _Spearman_ = 0.08, adjusted alpha level = 0.016; 0.05/3).

**Figure 3:**
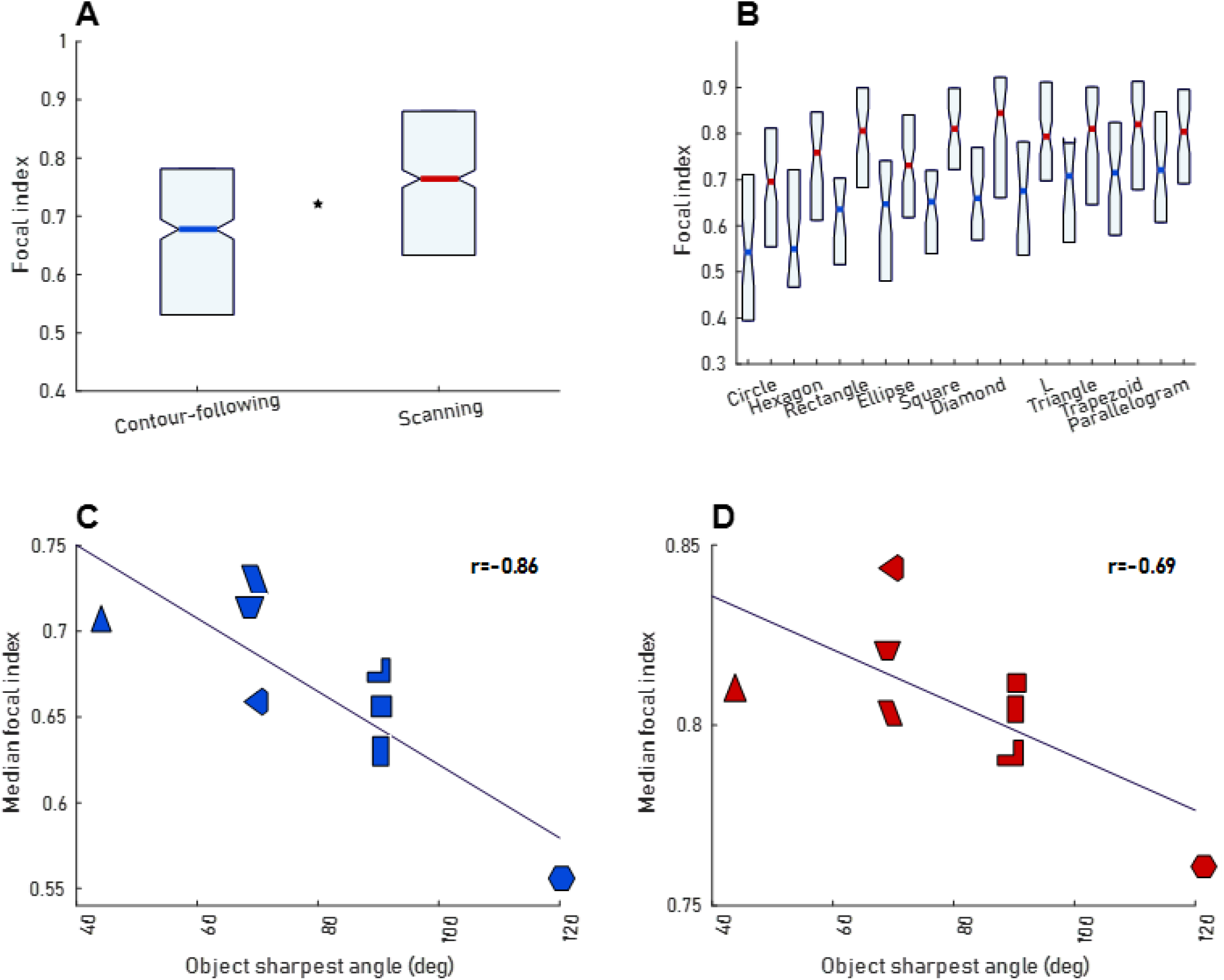
Focal palpation. **A** - The median of the focal index across all trials of all participants and all sessions (henceforth “grand median”) and quartiles for *CF* and *Sc* trials (p<0.005). Ns of subjects, sessions and *CF* and *Sc* trials and trials per subject are as in figure 2C. **B** - Grand median focal index per object in *Sc* and *CF* trials. **C**, **D** - Each data point represents the object grand median focal index and the object sharpest angle for either *CF* (C) or *Sc* (D) trials. N trials per shape, panels B-D: Triangle: 70 (Protocol I-55,III-15), Circle:128 (Protocol I-50,III-78),Rectangle:68 (Protocol I-60,III-8),Square −83 (Protocol I-49,III-34), Parallelogram-80 (Protocol I-54,III-26). Ellipse-121 (Protocol I-27, II-17, III-77), Hexagon: 66 (Protocol I-17, II-11, III-38). L: 104 (Protocol I-34, II-25, III-45). Trapezoid: 117 (Protocol I-61, II-35, III-21), Diamond: 95 (Protocol I-10, II-8, III-95).

### Palpation speed correlates with the participant’s spatial resolution

With adaptive active sensing it is expected that scanning velocities are tuned to optimize receptor activation. It has been shown that primates prefer hand movement patterns that preserve certain temporal cues (Cascio & Sathian, 2001; Drewing, 2012; Gamzu & Ahissar, 2001; Hollins & Risner, 2000; Lezkan & Drewing, 2018), cues whose temporal frequencies best fit one class of mechanoreceptors in the fingertip (Abraira & Ginty, 2013; Ahissar, 1998; Johansson et al., 1982; Talbot et al., 1968). Hence, adaptive active sensing predicts that variations in hand speeds across participants should correspond to variations in the spatial spacing of their receptors. In this case, the temporal frequencies of activation will be preserved in the preferred working range, as the temporal frequency generated on the skin when scanning a single edge is determined by the multiplication of the finger speed and the spatial frequency of receptors across the skin (Darian-Smith & Oke, 1980). To test for this possibility, we measured the Just-Noticeable-Difference (JND) in the index, middle and ring fingers of 10 of our participants (see Materials and Methods), as an indicator of the spatial spacing of the receptors array. Consistent with previous reports, JND values (JND = 3.16 ± 0.98) varied substantially across our participants (Peters et al., 2009). Importantly, for each participant the three tested fingers had similar JND values (Supplementary material, Fig. 4-Sp table 2, last column). During their first training session, the median tangential speeds of the participants were correlated with the mean JND measured across the three fingers (Supplementary material, Fig. 4-Sp10B, r = 0.74, p= 0.014, adjusted alpha = 0.0056; 0.05/9). The correlations were high for the JNDs of the middle (Fig. 4A, r= 0.84, p =0.0026; adjusted alpha level = 0.0045; 0.05/11) and index fingers and somewhat weaker for the ring finger (Supplementary material, Fig. 4-Sp10A,C, Index finger: r = 0.72, p = 0.017, Ring finger: r =0.42, p =0.2, adjusted alpha = 0.0056; 0.05/9). Participants were free to use any part of the three fingers array. Although we did not measure the movement of individual fingers, it is possible that the middle and index fingers were used more often than the ring finger, as previously reported in softness discrimination tasks (Katz, 2013; Zoeller & Drewing, 2020). A less frequent use may account for the weaker correlation of the ring finger. When crossing contours, the temporal frequency of activation of adjacent mechanoreceptors equals the scanning speed divided by receptor spacing (Darian-Smith & Oke, 1980). Assuming that receptor spacing equals the JND, the participant-specific mean frequencies ranged between 12.8 and 22.9 Hz (15.95 ± 3.18) during session 1. The positive correlation observed between the participants’ median speed and their JND was consistent with an attempt to maintain the temporal frequency within a narrow range (Carr, 1993). The mean evaluated activation frequency increased monotonically along consequent training sessions (Middle-finger: Ses1: 15.9 ± 3.18, Ses2: 17.07 ± 5.26, Ses3: 18.12 ± 4.9, Ses4: 18.14 ± 6.38) similar to the range of these activation frequencies (Ses1: 10.14, Ses2:19.2, Ses3: 15.45, Ses4: 22).

**Figure 4:**
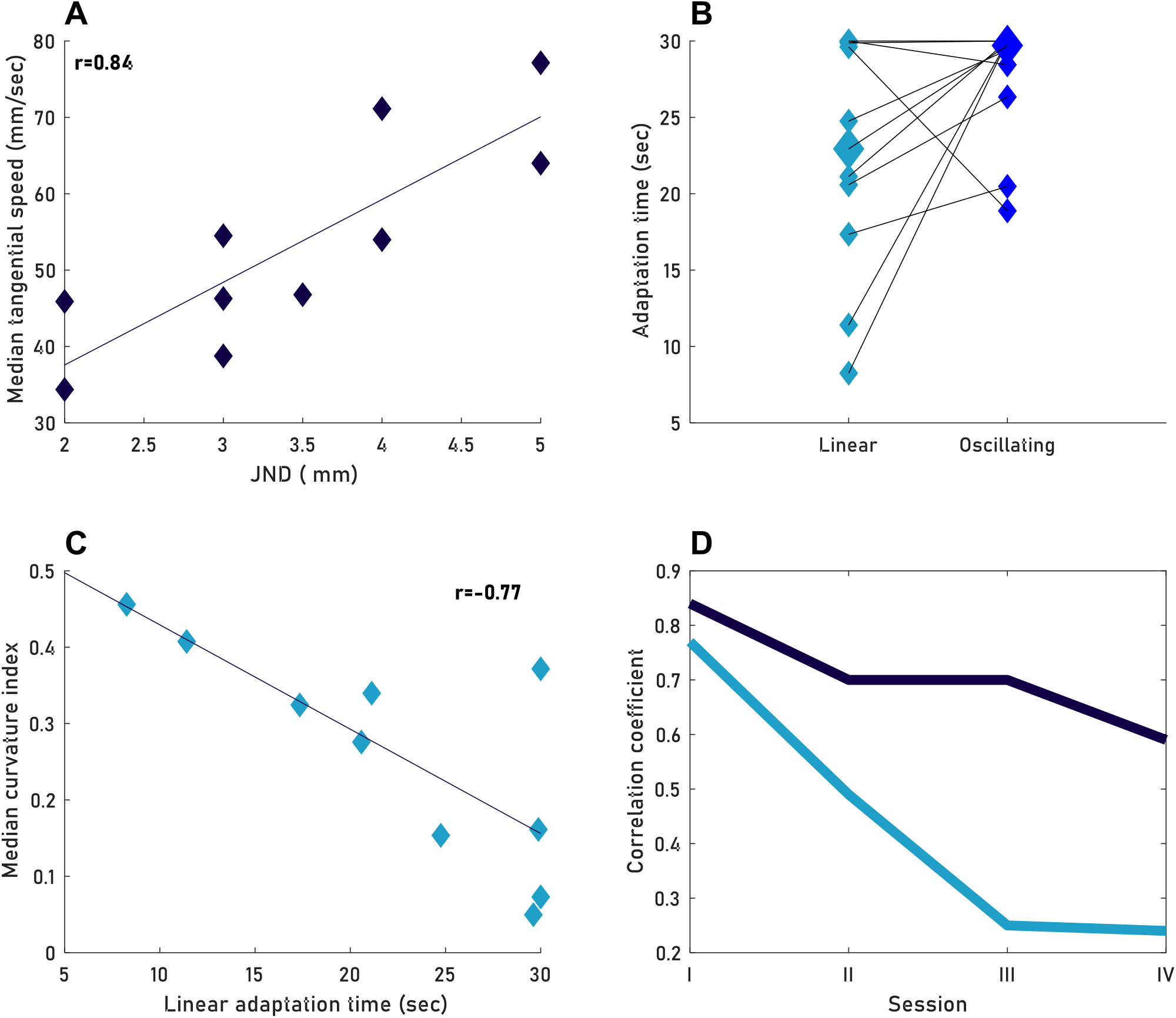
Dependency on sensory thresholds. **A** - Each data point represents the median tangential speed (across all trials of the first session) and the middle finger JND of one participant (18-29 trials per participant) **B** - Each two connected data points represent one participant mean adaptation time (Ta) using *Linear f*light blue*j* and *Oscillating (*blue) motion (11 *Linear* and 11 *Oscillating* trial per participant). Median values for each motion type are represented by a larger shape (median T_a *Oscillatory*_= 29.07, median τ_a *Linear*_= 22.9). **C** - Each data point represents the median curvature (across all trials of the first session) and the mean *Linear* τ_a_ of one participant (4-24 *CF* trials per participants/ **D** - Dark blue: Correlation coefficient between JND and tangential speed per session. Light blue: absolute value of the correlation coefficient between the curvature and adaptation time per session. N _participants_, all panels::10 (training protocol I-5, II-2, III-3).

The mechanoreceptor type that is most sensitive in the range of our evaluated activations frequencies in session 1 (between 12.8 and 22.9 Hz) is the rapidly adapting (RA) (Johansson et al., 1982). To test the potential effects of JND-dependent speed modulations on neuronal activations, we simulated the responses of grids of RA units with varying densities using a computational model (TouchSim; see Materials and Methods). The simulation shows that certain scanning speeds, for example the ones used by our participants in session one (between 34.4 to 77.14 mm/sec) generate activation rates within a narrow range (4 ± 0.05 spikes per edge crossing, using our simulation parameters; Supplementary material, Fig. 4-Sp11C-D). The simulation further shows that scanning speeds outside this range would generate larger variability in neuronal responses (std within speed rang = 0.05, std outside of speed range = 0.256, Supplementary material, Fig. 4-Sp11D).

### Palpation curvature inversely correlates with the participant’s sensory adaptation time

One of the major differences induced by the *Linear* and *Oscillating* sub-classes of *CF* (Fig. 2A) was in the duration of continuous interactions between the fingertip and the shape’s edge. *Linear* motion induced longer epochs of such interactions than *Oscillating* motion. A natural physiological feature that might be related to the choice between these strategies is the receptor adaptation rate – receptors with slower adaptation processes allow longer epochs of similar stimulation and longer *Linear* motions. We have thus tested the typical adaptation times of our participants, while scanning edges of different line types (straight, tilted or curved) or straight lines at different heights (see Materials and Methods). Participants were asked to follow outlines, forward and backward, for 30 sec. For each trial, the trajectory of the hand was analyzed and the time that elapsed from trial onset to the first deviation of the trajectory from the outline was considered as the adaptation time of that trial (Supplementary material, Fig. 1-Sp6A-B). The mean adaptation time of a participant (T_a_) was calculated across all her or his adaptation trials (see Materials and Methods). In order to examine if curvier *Oscillating* motion indeed elongated the effective adaptation time, we compared the adaptation times of our participants when following the shape’s outline using *Oscillating* and *Linear* motions (Experiment B). The adaptation times were longer with *Oscillating* motion for most of the participants (Mdn _*osculating*_ = 29.07 sec, IQR _*oscillating*_ = 30-26.3 sec vs Mdn _*Linear*_ = 22.90 sec, IOR _*Linear*_ = 29.8-17.3 sec; signed rank = 9, p = 0.12, Wilcoxon signed rank test; Fig. 4B).

To assess the pattern of *CF* quantitatively we computed the curvature index for every trial (Fig. 1D). During the first training session, the correlation between the participants’ τ_a_ (22.29 ± 8.02, N =10 participants) and their curvature index (0.26 ± 0.14) was high (Fig. 4C, r = −0.77, p = 0.0095, adjusted alpha = 0.01; 0.05/5). Thus, naïve participants with shorter adaptation times used curvier movements, such that they increased the number of border crossings and shortened the epochs in which skin stimulations remained relatively constant.

### The correlations with sensory threshold diminish with training

In the sessions that followed the first training session, hand speed and the curvature index gradually lost their dependence on the participant’s τa (Fig. 4D) or JND (Fig. 4D, supplementary material, Fig. 4-Sp10D). The decrease in correlation may result from transient, practice-induced changes in relevant sensory thresholds as previously reported (Wong et al., 2013), from strategy changes, or from both. One example of a strategy change is provided by two participants with short adaptation times – both abandoned the use of *CF* in favor of *Sc* (Supplementary material, Fig. 4-Sp12).

### Practicing on edge features increases tendency for palpating the shapes’ contours

During the first three practice sessions, eight participants palpated either the shapes of set A (Fig. 1A, black, training protocol I, 5 participants), or individual features (Fig. 1A, gray, training protocol II, 3 participants). During each of the practice sessions, *CF* palpation was significantly more common when palpating individual features (Fig. 5A, p<0.005 for all days). During the fourth testing session, all eight participants were tested on the shapes of set B (Fig. 1A, blue), which were presented to all participants in the same order. The difference in strategy choice in the practice sessions was preserved during this session (Fig. 5A, p<0.005) and was evident for each of the shapes (Fig. 5B, p<0.02 for all shapes, bootstrap p<0.05 for all shapes). Typical to *Sc* trials (Fig. 2C, Fig. 3A), the median focal index and the tangential speed were higher for those who practiced on shapes for the three preceding sessions (Fig. 5C, p<0.05). These participants were also faster in responding and showed higher confidence (Fig. 5C, p<0.02). These latter differences may result in part from the change in the form of the task for those who first practiced on individual features (from 3-AFC to 5-AFC). No experience-based difference was evident in recognition accuracy. The spatial distributions of mean visit rates during session 4 reflect the major palpation difference – participants who first practiced with individual features (Fig. 5D, bottom row) focused more on the borders compared with those first practicing on shapes (Fig. 5D, top row).

**Figure 5:**
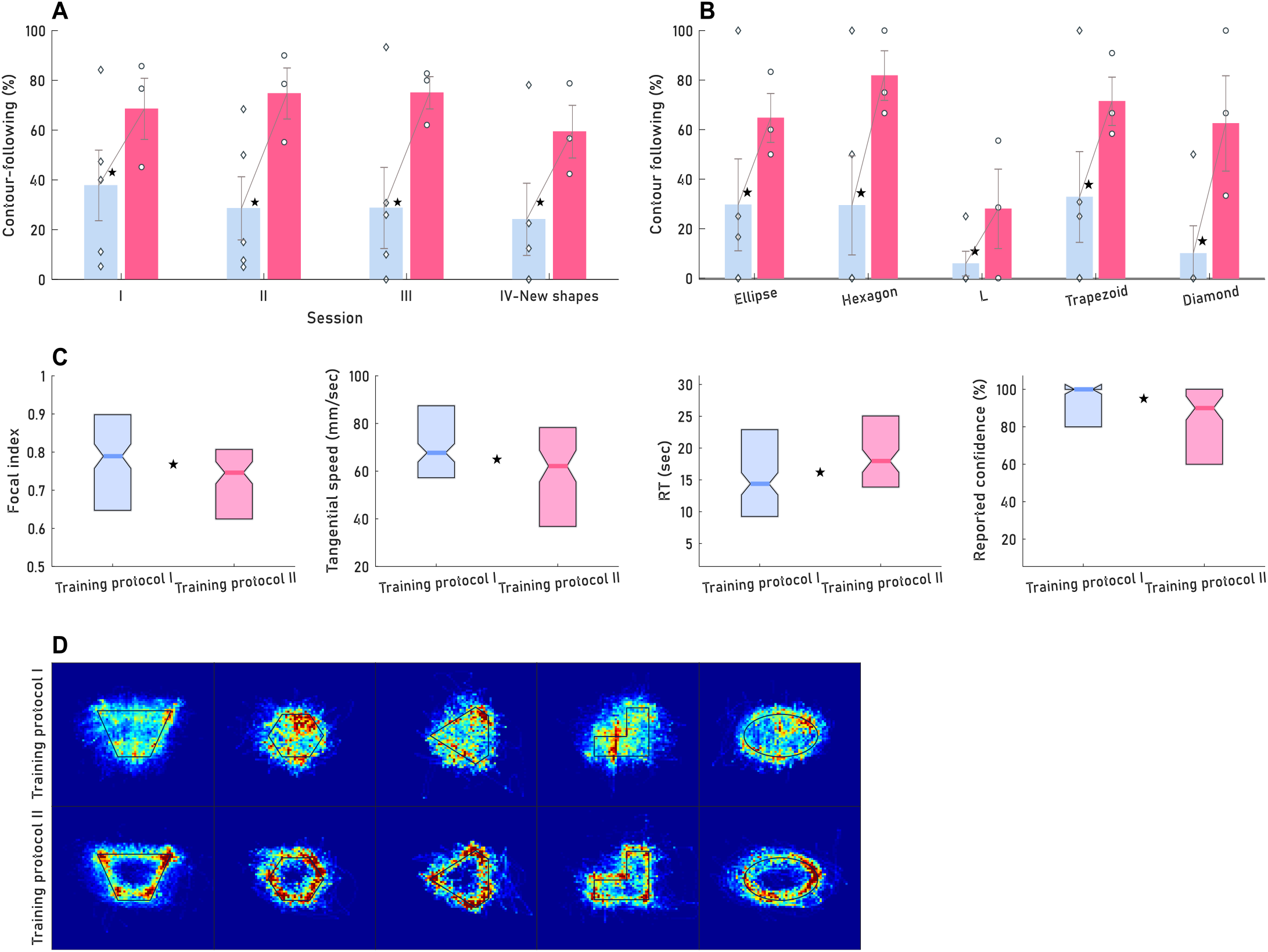
Dependency on practice history. **A, B** - Prevalence of contour following trials for training protocol I (light blue) and II (pink). **A-**For each practice session. N _trials Sessions I-III_: Protocol I: 84-94 trials per session, 15-20 trials per subject in each session. Protocol II: 87-89 trials per session,28-30 trials per each subject session. N _trials Sessions IV:_ protocol I-149 trials (24-32 trials per subject), protocol II: 96 trials (30-33 trials per subject). **B**-For each shape at the 4^th^ practice session. N _Trials per shape_: Ellipse-44 (Protocol I-27, II-17), Hexagon:38 (Protocol I-17, II-11). L: 59 (Protocol I-34, II-25). Trapezoid: 96 (Protocol I-61, II-35), Diamond: 18 (Protocol I-10, II-8).Training protocol II used significantly more *CF* in all sessions (χ^2^ _Session I_ (1, N=179) =17, χ^2^ _Session II_ (1, N=171) =36.45, χ^2^ _Session III_ (1, N=182) =38.95, χ^2^ _Session IV_, (1, N=245) =30.94) and for all shapes (χ^2^ _Ellipse_ (1, N= 44) =5.23, χ^2^ _Hexagon_ (1, N=28) =7.33, χ^2^_L_ (1, N= 59) =5.45, χ^2^_Trapezoid_(1, N=96) = 13.33 χ^2^ _Diamond_ (1,N=18) =5.51. In panels A-B, error bars represent the standard error of the mean, each data point represents the prevalence of *CF* trials for one subject. **C**-Median and quartiles of focal index and tangential speed, RT and reported confidence for training protocol I and II (p<0.05 for all differences). **D**-Visit rates of all participants of the shapes practice training protocol (training protocol I, top) and those of the features practice training protocol (training protocol II, bottom) during session 4. N _subjects_, all panels: =8 (Protocol I-5, III-3).

## DISCUSSION

This paper describes the repertoire of motor strategies employed by human participants when required to recognize planar (2D) object shapes. Consistent with previous studies of 3D shapes (Klatzky & Balakrishnan, 1991; Lederman & Balakrishnan, 1991; Lederman & Klatzky, 1987), we found that one major strategy was Contour-Following (*CF*) – participants moved their fingers along the contours of the shape. In addition, we found a second major strategy – Scanning (*Sc*) – with which participants used large movements crossing the entire shape. Both strategies, and in particular *Sc*, included focal scanning of specific object regions. *CF* was employed in two major patterns – *Linear* and *Oscillating* motion.

While none of our physiological measures could fully predict participants’ choice of palpation strategies, they could account for a significant portion of their variability. The naïve tendency to perform an *Oscillating* motion variant of *CF* was correlated with having faster mechanosensory adaptation. In addition, at session one the tangential velocities were correlated with the spatial resolution of mechanoreceptors in the fingers, as indicated by the participants’ JNDs, such that the temporal frequencies generated by the interaction with the shapes’ contour were between 13 to 23 Hz. These correlations were high when our participants were naïve to the task and gradually decreased with practice. One interpretation of these results is that, with practice, human participants adapt scanning strategies that compensate for their sensory limitations.

Taken together, these results open a window to the complex ways in which individual sensory abilities, practice repertoire and task-demands converge to an effective motor-sensory exploration strategy.

### Experimental constraints

In these experiments we attached the three palpating fingers (index, middle and ring) together, in order to be able to measure all motion components that are relevant to the perception of the objects. The participants were free to use any part of the affective sensory array, which was composed of the combined contact areas of the three fingertips. In a way, the three fingers were treated as one large finger. As far as known from the literature, tactile perception naturally involves coordinated motion of all contacting fingers: Participants’ fingers were shown to have correlated positions and speeds in tactile search tasks (Morash et al., 2013) and in a task designed to require only one finger (Fish & Soechting, 1992). When human participants were specifically instructed to use only one finger, low amplitude, correlated movements were seen in the other fingers (Hager-Ross & Schieber, 2000). Nevertheless, in the current study we did not examine the independent movement of each finger, a degree of freedom that might be treated differently by different participants.

### Exploratory procedures in 3D and 2D objects

In a pioneering study, Lederman and Klatzky termed the concept ‘‘Exploratory Procedures” (EPs) to refer to the motor-sensory exploration strategies characterizing the palpation of various classes of 3D objects under specific tasks (Lederman & Klatzky, 1987). Here we zoomed into a one class of objects – planar shapes – and tried to characterize the EPs that are naturally employed by human participants when asked to identify shapes. Our results expand those of Lederman and Klatzky and their collaborators (Klatzky & Balakrishnan, 1991; Lederman & Balakrishnan, 1991; Lederman & Klatzky, 1987) by showing that people employ not only *CF*, as observed by these researchers, but also *Sc* motions, when exploring shape in 2D objects (Fig. 2). The *Sc* motion resembles the ‘Lateral-motion’ EP, which was reported as the dominant strategy for texture palpation in 3D objects (Lederman & Klatzky, 1987). Our high-resolution tracking system provided additional information about the nature of these strategies, revealing that human participants vary in the specific pattern of each EP – *Linear* and *Oscillating* motions for *CF* and uniform and non-uniform motions for both the *Sc* and *CF*. Planar shape palpation also revealed a frequent use of a palpation policy that was not describe before – focal palpation. Participants often dedicated a significant portion of time to explore specific regions on the objects. Focal palpation was more evident in *Sc* trials, (Fig. 3) and was depended on the properties of the explored object: Objects with sharp angles were explored in a more focal manner. The tendency to use focal palpation was hardly affected by the participants’ practice history, nor by their physiological thresholds.

### Dependency of palpation strategies on practice history

Numerous behavioral studies show that tactile perception depends on practice history (Kaas et al., 2013; Sathian & Zangaladze, 1997; Spengler et al., 1997; Withagen et al., 2013; Wong et al., 2013). Here we focused on the choice of palpation strategies and its dependency on practicing on shapes versus practicing on individual features. The findings were clear: Practicing on individual features entails a strong preference for using the *CF* strategy. This preference was preserved during an entire testing session of shape palpation that followed three practice sessions of feature palpation (Fig. 5).

### Sensation-dependent movements, controlled variables and closed-loop touch

Sensory-motor behavior depends on the physiological parameters of sensory receptors (Neubarth et al., 2020; Severson et al., 2017). Consistently, our results clearly demonstrate that the choice of motion strategy depends strongly on sensory physiology. The speed of motion decreased with increased spatial resolution (reduced JND) at the fingertip, and the degree of motion curvature increased with faster adaptation times (Fig. 4). These correlations were very strong when our participants were naïve to the task: During session 1, the physiological measures, JND and Ta, could explain 70% and 59% of the variability in the kinematic measures, tangential speed and curvature index, respectively. Such behavior is expected when the tactile system is concerned with maintaining specific sensory variables in their “working ranges”, i.e., ranges that allow satisfactory perception (Ahissar, 1998; Ahissar & Vaadia, 1990; Marken, 2001). Thus, rats maintain head azimuth and whisker speed (Saraf-Sinik et al., 2015) and humans maintain hand coordination and hand speed when localizing objects around them (Saig et al., 2012). When scanning planar objects, humans often attempt to maintain temporal activation variables within certain ranges, as well. Thus, humans reduce hand speeds with higher external spatial frequencies when perceiving textures (Gamzu & Ahissar, 2001) and modify radial and tangential forces, together with lateral hand speeds, when exploring surfaces with different geometry and friction attributes, such as to maintain a certain amount of skin deformations (Smith et al., 2002). Maintaining the sensory signals within certain ranges is expected to facilitate their predictive processing within brain circuits (Ahissar & Vaadia, 1990; Barascud et al., 2016; Huang & Luo, 2019; Rao & Ballard, 1999). Variables that are actively maintained within specific ranges are termed “controlled variables” (Biswas et al., 2018; Burstedt et al., 1997; Marken, 2001; Todorov & Jordan, 2002). Preserving them in preferred working ranges requires a closed-loop architecture, in which the variables can be sensed and manipulated. As these controlled variables are serving perception, their control is likely to optimize sensation (Ahissar & Assa, 2016; Biswas et al., 2018; Buckley & Toyoizumi, 2018b; Powers, 1973; Simony et al., 2008; Thomson, 1927).

Controlling hand speed in the current experiment resulted in maintaining the mean temporal frequency of fingertip activations, when crossing contour edges, between ~10 to 40 Hz across participants and sessions. In texture-related tasks the effective temporal frequency of individual receptor activation is typically maintained between 15 – 30 Hz (Gamzu & Ahissar, 2001). This frequency range is optimal for activating rapidly-adapting (RA) receptors at the primate fingertip (Ahissar, 1998; Johansson et al., 1982; Talbot et al., 1968). Emphasis of RA receptors in this experiment is in line with the fact that the height of our shapes was 25 microns, which is better sensed by RA receptors compared with SA receptors (Johansson et al., 1982; Neubarth et al., 2020; Talbot et al., 1968).

Our simulations further suggested that the JND-dependent control of scanning speed observed here (Fig. 4A) resulted in a quite uniform activation rate of RA receptors (Supplementary material, Fig. 4-Sp11C-D). Another support for the conjecture that the tactile system attempts to maintain uniform activation rates comes from our behavioral adaptation measurements. These results suggest that our participants made an effort to prevent receptor adaptation - participants with shorter adaptation times used curvier movements (Fig. 4C), which were likely to reduce receptor adaptation and thus maintain activation levels.

### Conclusion

Our results suggest that a variety of tactile palpation strategies are used for shape recognition and that the choice of strategy and palpation parameters are affected by the object’s spatial parameters as well as by the individual sensory physiology and practice. All these variables interact in the control of scanning movement. While the results reported here do not sum up to a complete model of tactile perception of shapes, they provide an initial framework detailing the interactions of several key variables in this process.

## MATERIALS AND METHODS

### Overview of experimental design

This study was composed of two experiments. The first experiment (Experiment A) was designed to study the characteristics of tactile scanning when perceiving planar shapes. In the second experiment (Experiment B, conducted 18 months after Experiment A), we measured adaptation times and spatial resolutions of most (10 out of 11) of the participants who took part in Experiment A. The experimental procedures were approved by the Helsinki committee of the Tel Aviv Sourasky Medical Center.

### Experiment A: Shape recognition

#### Participants

Eleven right-handed participants [seven female and four male students, aged 21-32, (25.36 ± 3.17) years] took part in the study. Informed consents were obtained from all participants, in accordance with the approved declaration of Helsinki for this project. The participants were paid for their participation. None of the participants had any previous experience with tactile recognition tasks.

#### Experiment design and procedure

All training protocols followed the same basic procedure. Participants were asked to identify two-dimensional (2D) engraved shapes or features (Fig. 1A). Before the beginning of the first session, participants saw illustrations of all the shapes or features that were presented in this session, and they were shortly trained by palpating on one of the shapes or features until they reported that they understood the task. Participants performed four or five experimental sessions. Whenever new shapes were introduced, a visual illustration of them was presented to participants at the start of the relevant session. At the beginning of each trial, the experimenter placed the participant’s gloved-hand (Fig. 1B) a few centimeters above the center of the shape or feature board, and the trial began when the participant was allowed to put her or his hand on the shape (see ‘Testing apparatus’ below). Participants were requested to raise their palpating hand and name the shape placed in front of them when they identified it, as well as to report their confidence level. Trials ended with the participant’s declaration or after a time limit of 30 seconds (sec) (whichever came first). A feedback was given on whether the answer was correct or not. In each session or block (see below), the order of trials was randomized and corrected such that each object was presented at least once. Trial presentation order within a session or block was kept constant across participants.

Three different training protocols were used (Supplementary material, Fig.1-Sp1) in order to allow protocol independent observations:

*Training protocol I - Shapes*: Five participants were trained for three sessions on a fixed set of geometrical shapes (Fig. 1A, Set A, black, i.e. triangle, circle, square, etc.) in a five-alternative forced choice (5-AFC) recognition task. Each session included 20 trials. The order of trials was random but kept constant between participants.
*Training protocol II-Features*: Three participants were trained for three sessions on the features set (Fig. 1A, gray). Features were presented in three blocks, 10 trials each: ‘Angle’ block (ninety, acute or obtuse angles), ‘Tilt’ block (tilt-right, tilt-left or straight vertical line) or ‘Curvature’ block (concave, convex or straight vertical line). Before each block, participants were notified which block is presented, and they were requested to name the presented feature in a three-alternative forced choice (3-AFC) recognition task. The order of blocks was randomized between participants and kept constant per each participant in all three sessions. The order of trials within each block was randomized but kept constant between participants.
*Training protocol III - Non fixed shapes*: Three participants practiced for five sessions. Each session included 35 trials. In the first session, the stimulus set included five shapes (Fig. 1A, ellipse, circle, rectangle, parallelogram, square). In each of the following sessions, at least one shape was removed and instead, a new shape was introduced (Supplementary material, Fig.1-Sp1, right, yellow). A visual illustration of this new shape was presented at the beginning of the session. Although only five shapes were presented in each session, participants were not told that one of the previously presented shapes will not appear. Thus, the task gradually changed from a 5-AFC discrimination task (session one) to a 9-AFC recognition task (session five). The objects presented in each session were: <u>Session one</u>: Ellipse, circle, rectangle, parallelogram, square. <u>Session two</u>: Ellipse, circle, *diamond*, parallelogram, square. <u>Session three</u>: Ellipse, circle, diamond, parallelogram, *L-shape* (L). <u>Session four</u>: Ellipse, circle, diamond, *hexagon*, L. <u>Session five</u>: *Triangle, trapezoid*, diamond, hexagon, L.

##### Testing session

Participants who trained in protocols I and II were tested during their 4^th^ session on a set of novel geometrical shapes (5-AFC, Fig. 1, Set B, blue). This session included 34 trials.

#### Hand tracking

Hand motion was tracked in 3D coordinates (x, y, z), using Vicon 612 motion capture system (Vicon Motion Systems Ltd, Oxford, .UK) and the Nexus 2.5 software. A custom ‘Vicon labeling Skeleton Template’ (VST) of the hand was designed. The VST included three segments and four markers attached above the wrist carpal bones and the middle finger’s middle and proximal phalanges (Fig. 1B). At the beginning of each session, the VST was calibrated for the current subject’s parameters, and a labeling skeleton file (VSK) was created. Hand motion was sampled at 200 Hz (76.5% of trials), 100 Hz (5.7%) and 240 Hz (17.8%), (see ‘Vicon tracking’ within ‘Data Analysis’ below).

#### Tactile objects

Objects were engraved on aluminum boards of size 150X150 mm, such that the area inside the shape was raised to 25 μm in relation to the board surrounding. Objects were divided into three sets: Shapes – sets A & B (hereafter referred to as ‘shapes’, Fig. 1A, black and blue) and Features set (Fig. 1A, gray).

#### Testing apparatus

Experiment took place in a Vicon arena. Participants sat in front of a table on which the stimulus was placed. The aluminum board was placed in a plastic frame, preventing its movement. Participants were blindfolded and wore a glove on their right hand. Four Vicon designated markers were connected to the glove, in a way that two markers were connected above the middle and proximal phalanges of the middle finger and two above the wrist carpal bones (Fig. 1B). In order to simplify and standardize the experiment, we reduced the degrees of freedom by banding together the index, middle and ring fingers with a tape-as if all three fingers belong to one surface plane. This restriction facilitated analysis and allowed comparison to other existing datasets. Three corresponding fingertips of the glove were cut, such that the finger pads were uncovered. Two glove sizes were used and chosen according to participant’s hand size.

#### Data analysis

##### Vicon tracking

Trajectories of one marker were analyzed – the ‘tip’ marker that was placed above the middle finger, middle phalanges (closest to the finger pad, Fig. 1B). In a fraction of the sessions, the number of performed trials was smaller than planned (10 out of 47 sessions, 21.27%). A fraction of performed trials was excluded due to missing capture frames; the analysis throughout the paper includes the remaining number of trials (1196 out of 1367 trials, 87.4%). A fraction of these trials was not sampled at 200 Hz but at 100 Hz (5.7%) and 240 Hz (17.8%). These trials were resampled to 200 Hz: 100 Hz trials were linearly interpolated by a factor of two using the MATLAB function ‘interp1’; 240 Hz trials were linearly interpolated by a factor of five using the ‘interp1 ‘MATLAB function and then down sampled by a factor of six using the ‘downsample’ MATLAB function. In some sessions (n = 22 out of 47 sessions, 46.8%), the calibration of hand position in relation to the shape was lost. To compensate for differences in calibration quality, we chose for each session one trial with clear hand-trajectory orientation relative to the borderlines of the shape and calculated the translation and rotation between this trajectory and the shape. We used these parameters to shift and rotate all the hand trajectories in that session.

##### Trial start, end and reaction time

At the beginning and end of each trial, participants lowered or raised their hands, respectively (see experimental procedure). The point of hand lowering was marked as the first frame in which the height in the z-axis was equal or smaller than the median z height in the rest of the trial. The point of hand raising was marked in a similar way: We smoothed hand velocities in the z-axis (using a moving average, MATLAB function, ‘movmean’, window size = 30 samples). A histogram of all data was plotted, forming a bi-modal distribution (Supplementary material, Fig. 1-Sp2). Hand raising was marked as the first frame in which z-speed was higher than a threshold marking the second mode. In addition, in order to exclude 2D movements that accompanied hand lowering or raising, trial start (t = 0) was defined as the first frame after hand lowering in which the hand was at <5 mm from the shape outline. Trial end was marked as the last frame for which all consequent frames were >5 mm from the contour. Trial reaction time (RT) was the difference between trial start and trial end.

##### Classification of movement types

Following initial screening of trial trajectories, we aimed at classifying the trials into two types: Contour-Following (*CF*) trials were defined as trials in which the hand remained in the vicinity of the object outline. Scanning (*Sc*) trials were defined as trials in which the trajectory crossed between distant parts of the objects’ contour. Trials were classified according to this distinction using two methods: Algorithmic and perceptual (by human observers). The analysis throughout the paper is based on the algorithmic classification. The human observer classification was used as a verification method. The analyses based on it are presented as supplementary material (Supplementary material, Fig. 1-Sp4).

###### Algorithmic classification

The algorithm used heuristic criteria that were based on our initial inspections of motion strategies. For each original shape two similar “guiding shapes” were plotted around its centroid: An inner and outer guiding shapes whose areas were 0.25 and 1.5 of the original shape, respectively. The ‘L’ and ‘convex’ shapes were exceptional (see Supplementary material, Fig. 1-Sp3). Scanning trials (*Sc*) were trials in which either the hand crossed the inner guiding shape (Fig. 1C, left), or those in which the hand did not remain between the inner and outer guiding shapes for more than 75% of the time (Fig. 1C, second left), or both (Fig. 1C, second right). *CF* trials (Fig. 1C, right) were trials in which the hand remained in the buffer between the inner and outer guiding shapes for more than 75% of the time, and did not cross the inner guiding shape

###### Human observer classification

Two human observers classified the trials – one of the authors (NM) and a naïve observer (SM). Each observer classified all trials twice consecutively, and the second set of labeling was used for analysis. The observers could classify a trial as either *CF*, *Sc* or *Oth*er. The two observers differed in the percentage of labeling trials as ‘*Other*’ (NM, 11.8%; SM, 15.6%) and agreed on 66.6% of the trials (78% when *Other* trials were excluded).

##### Trajectory curvature

Curvature evaluation was calculated in the following way: Each trial’s hand trace was smoothed using a moving average with a window size of three samples. The trace was divided into segments of 20 mm length; the overlap between consequent segments was 0.01% (0.2 mm). Curvature index was defined as the subtraction of the shortest distance between the beginning and end of the segment’s coordinates from the segment-traveled distance, divided by the traveled distance:

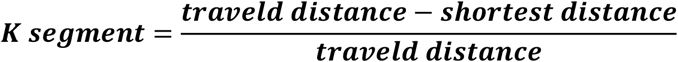

The value of the curvature index is in the range between 0 to 1: The index is closer to zero as segment trace resembles a straight line (Fig. 1D, dark blue) and to one as it is more curved (Fig. 1D, dark red). The median of the indices of all segments in a trial was assigned as the trial curvature index. The constant-length segmentation was preferred over the constant-time segmentation because stronger curvatures are usually accompanied by slower movement (Viviani & Flash, 1995; Viviani & Schneider, 1991), which would lead to over-estimation of curved movements in the latter case.

##### Focal index

To evaluate the distribution of palpation density along the shapes’ outlines, a focal index was used. Circles at a radius of 10 mm were plotted on the object outline, such that the distance between their centers was 0.5 mm. The traveled hand-trajectory distance in each circle was computed. As a measure of dispersion, the difference in traveled distance between the most visited (Fig. 1E, left, purple circle) and least visited (Fig. 1E, left, green circle) areas was computed (Fig. 1E, right) and divided by their sum.

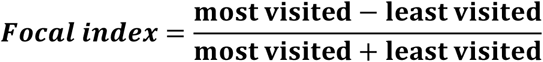

The least-visited circle was determined after removing circles that were not visited or circles that overlapped with non-visited circles (Supplementary material, Fig. 1-Sp5).

##### Trial entropy

Was calculated using the MATLAB function ‘entropy’, calculating the entropy of a gray scale image. A gray scale image of each trial was created using the MATLAB function ‘hist3’. The entire stimulus board (150×150 mm) was included. Bins with the size of 2×2 mm were used.

##### Visit rates

The number of visits of the hand’s trajectory in each 2×2 mm bin were counted and divided by trial duration.

### Experiment B: Spatial and temporal thresholds

In this experiment, we tested the spatial resolution and temporal adaptation profiles of 10 of our participants.

#### Participants

Ten participants who participated in experiment-A took part in this experiment. The participants were naïve to the purpose of the experiment and were paid for their participation. Informed consents were obtained from all participants, in accordance with the approved Declaration of Helsinki for this project.

##### Task 1. Temporal profile of sensory adaptation

###### Motion tracking

Trials were filmed (sampling frequency – 30 Hz) and were later analyzed using the MATLAB vision toolbox and the ‘tracker’ function.

###### Tactile objects

The ‘straight’, ‘right tilted’ and ‘convex’ outlines from the features set were used (Fig. 1A, right, gray). In addition, we used eight identical-size rectangles raised to different heights, which were engraved on an aluminum board. Rectangle heights ranged from 10 μm to 80 μm and differed from one another by 10 μm.

###### Testing apparatus

Apparatus was identical to the apparatus in Experiment A, apart from the fact that it did not take place in the Vicon arena. The same glove was used, which had one polyester marker connected above the middle phalanges of the middle finger.

###### Design and procedure

This experiment aimed to test the temporal profile of sensory adaptation while following the contour of our tactile objects. The participants were requested to use two types of motions: *Linear* and *Oscillating* (Fig. 2A). At the beginning of the session, these motions were demonstrated by the experimenter and then practiced by the participants, first on a desk and later using the ‘rectangle’ stimulus board (Fig. 1A, Set A, black). Before each trial, participants were instructed which motion type they should use. After each trial, a break of 45 sec was taken, in which participant’s hand was placed such that the finger pads were in the air. This was meant to allow full recovery of both slowly and rapidly adapting receptors (Leung et al., 2005). Before each trial, the participant’s hand was placed on the starting point of different outlines, and they were asked to report if they could feel the contour. Participants were instructed to follow the contour using *Oscillating* or *Linear* motion until the experimenter told them that the trial ended. Trial duration was 30 sec. During each trial, the contour was tracked either once or more (moving back and forth along it), depending on hand velocity. The participants were not asked to report anything nor were they given any feedback.

###### Stimuli Types

Two tactile arrays were used:

<u>*Array with different outline types:*</u> The outline was either ‘straight’, ‘right tilted’ or a ‘convex’ engraved shapes raised to 25 μm (Fig. 1A, Features set, gray). This task included six trials, as each outline type was followed using both motion types (*Oscillating* and *Linear*). The order of the trials was random and kept constant between participants.
<u>*Array with different outline heights:*</u> A board with 8 rectangles was used. Rectangle lines heights ranged from 10 μm to 80 μm and differed from one another by 10 μm. The task included 16 trials, as each outline height was followed using both motion types. The task was performed in two blocks - *Oscillating* and *Linear*. The order of blocks was random and kept constant across participants. The order of trials within each block was randomized but kept constant across participants.

###### Data analysis

For each trial we identified the first time in which the tracking hand lost the outline using visual inspection of the trial’s movies. Two human observers examined the movies – one of the authors (NM) and a naïve observer (SG). Each observer marked the first point in which the hand’s trajectory clearly deviated from the outline and assigned the time duration between trial start and the point of deviation as the trial T_a_ (Supplementary material, Fig. 1-Sp6A-B). In trials in which participants did not deviate from the outline the observers assigned T_a_ as the maximal trial duration (30 sec, Supplementary material, Fig. 1-Sp6C-D). Trials in which it was not clear whether participants indeed lost the outline were excluded. For the majority of trials (n = 170 out of 225, 75.56%) both observers confidently assigned a T_a_. For these trials, the correlations between the T_a_s assigned by the two observers, for *Oscillating* and *Linear* motion trials, were r = 0.73 and r = 0.68, respectively (p<0.005, Supplementary material, Fig. 1-Sp7A-B). The distribution of the differences between the two observers’ T_a_s exhibited a clear mode at 0 and a secondary mode between 0 to 5 sec (Supplementary material, Fig. 1-Sp7C). In this analysis we included only trials for which the difference was < 5 sec, which composed 86.47% (147 out of 170) of the trials. For these trials, the T_a_ was taken as the mean of both observers’ T_a_s.

##### Task 2. Spatial resolution

The spatial just-noticeable difference (JND) of each participant was measured using a static two-point discrimination test, applied to the pads of index, middle and ring fingers, similar to a procedure previously described (Louis et al., 1984). Participants placed their hand comfortably on a table and were blindfolded. Participants were asked to report whether they feel contact in one or two points on their skin. The task was demonstrated on the participants’ forearm before starting the experiment. An adjustable compass was used. The interval between the two tips of the compass was gradually reduced until the participant could not differentiate between the two points. An effort was made by the experimenter to keep the same amount of pressure. Threshold was determined as the first interval at which the two points could not be distinguished. The order of measured fingers was ring, middle and index finger for all participants.

### Statistical analysis - experiments A & B

Unless stated otherwise, the compared distributions were tested for normality using the Anderson-Darling test. If at least one of the compared distributions was recognized as non-normal, the Mann–Whitney U test (non-paired comparisons, two tailed), or Wilcoxon signed-rank test (paired comparisons, two tailed) were used. Otherwise, a two-tailed (independent or paired) t-test was used. Categorical data was tested using the Chi-square test of independence using all of the trials in each training protocol session (84-146 trials). Multiple comparisons were corrected using the Bonferroni method. Since in some of the compared distribution part of the observations were dependent (trials belonging to the same subject), significance was additionally tested using bootstrap. Unless stated otherwise, all significance reports are based on both a model-based (one of the aforementioned) and a bootstrap test.

Bootstrap was performed in the following way: Trials from both of the compared groups were mixed to one pool. Ten thousand iterations were used. At each iteration, two samples were taken from the mixed pool. One sample was at the size of the number of trials of comparison group I and the second at the size of the number of trials of group II. A difference calculation was repeated for each iteration. The fraction of bootstrap values that were more extreme (larger or smaller, depending on the case) than the experimental one was reported as the boundary of the probability of getting the experimental value by chance.

### Simulations

To test the effects of JND-dependent speed modulations on neuronal activations, simulations of mechanoreceptor responses were performed using TouchSim (Saal et al., 2017). Seven grids of rapidly-adapting (RA) mechanoreceptive units were formed with inter-receptor distances of 1 to 4 mm, with 0.5 mm intervals. The resulting grid sizes were between 10.5 x 10.5 and 12.5 x 12.5 mm (Supplementary material, Fig. 4-Sp10A). The stimulus was comprised of two columns of pins, each with a radius of 0.5 mm, aligned between the two left-most receptor columns (Supplementary material, Fig. 4-Sp10B). The pin radius was 0.5 mm. In each simulation run, the two stimulus columns were pressed onto the finger grid with a delay that corresponded to the simulated scanning speed. Scanning speed was varied between 20 to 250 mm/sec (with intervals of 10 mm/sec) and pressing duration was 0.2 sec. The duration of each simulation was 1 sec and the sampling rate was 5000 Hz. The resulting spike counts were smoothed (moving average, window size = 5 samples).

## Acknowledgements

We thank Nir Knaan and Yahel Zamosh for helping in designing and conducting experiment A and Silman Bensmaia for insightful comments and advices. This research received funding from the European Research Council (ERC) under the EU Horizon 2020 Research and Innovation Programme (grant agreement No 786949). E.A. holds the Helen Diller Family Professorial Chair of Neurobiology. A.A. holds the Sam and Frances Belzberg Chair in Memory and Learning.

## SUPPLAMENTRY MATERIAL

**Figure 1-Sp1:**
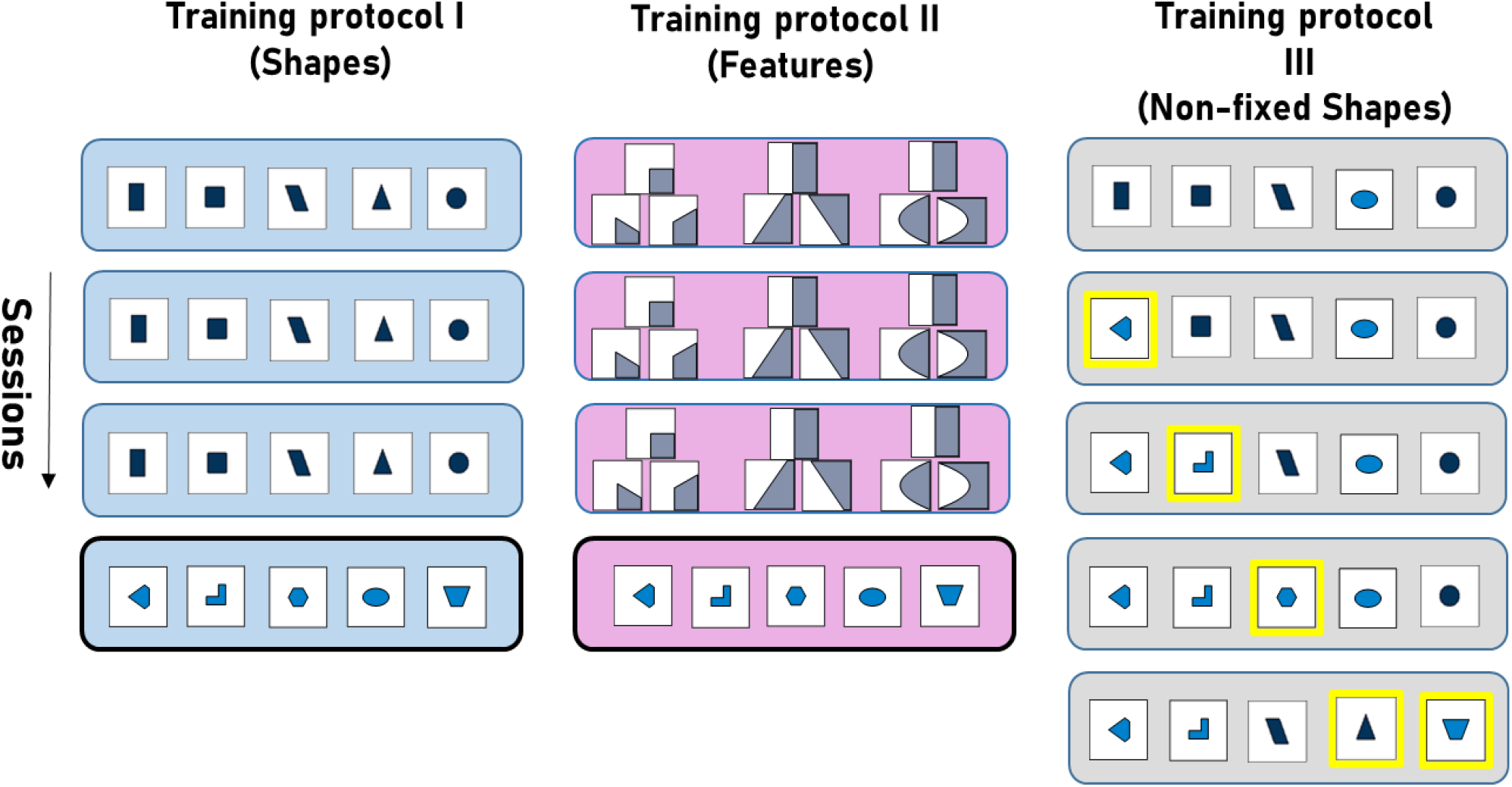
Training protocols: Training protocol I (left, blue, Shapes) practiced on set A shapes (black) for three sessions, and on set B (blue) at session four. Training protocol II (middle, pink, Features) practiced on the features set (gray) at the first three practice sessions. Features were presented in three blocks: Angle (left, ninety, acute or obtuse angles), Tilt (middle, tilt-right, tilt-left or straight vertical line) and Curvature (right, concave, convex or straight vertical line). Training protocol II then practiced on set B (pink) at session four. Training protocol III (right, gray, Non-fixed shapes) practiced on shapes from both sets A and B. Starting from session two, at least one new shape (yellow) replaced a previously presented shape.

**Figure 1-Sp2:**
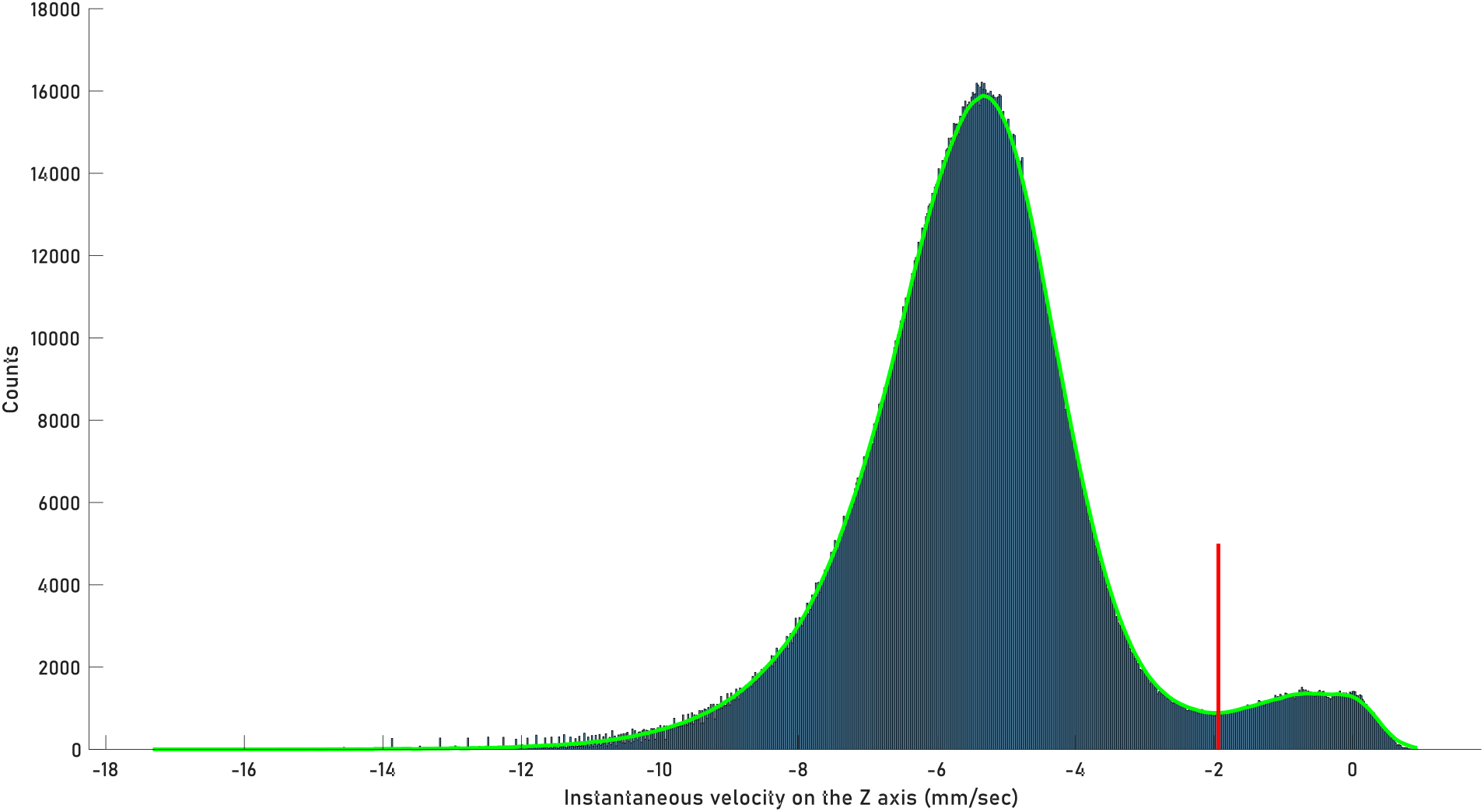
Distribution of hand instantaneous velocity speed (including all trials) at the z-axis: The curve is smoothed using a moving average (MATLAB function, ‘movmean’, window size = 30 samples). A threshold marking the second mode is plotted (red). A single trial’s end was marked as the first frame in which z-velocity was higher than this value.

**Figure 1-Sp3:**
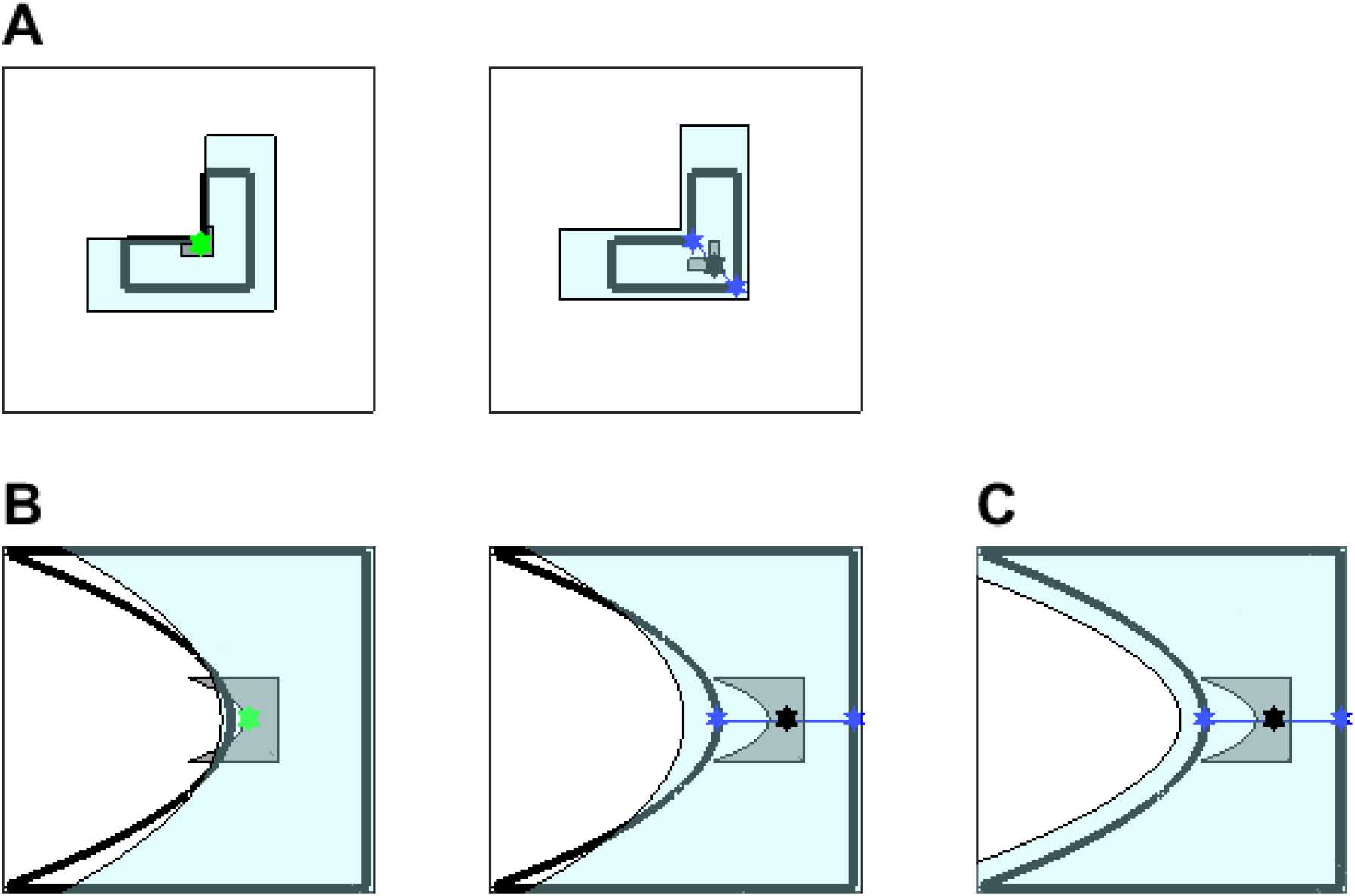
Classification of L and convex trials: To allow for trial classification, a smaller area object (0.25 smaller, gray) and a larger area object (1.5 larger, light blue) were plotted around objects (black line) centroid (green). In the case of the ‘L’ (A, left) and ‘Convex’ (B, left) the centroid (green) lies close to the object contour. Therefore, different points within the object were chosen: For the ‘L’ a middle point (black) between two points on the contour (blue) was chosen, and the small and large objects were plotted around this point (A, right). For the ‘Convex’, similarly, a middle point (black) between the blue marked points was chosen (B, middle) and the small and large object were plotted around this point. For the ‘Convex’ feature, this definition still did not suffice for the larger object to completely contain the original shape (B, middle). Therefore, the larger shape was finally obtained by replotting the shape’s border at a distance increased by 10 mm in the relevant direction

**Figure 1-Sp4:**
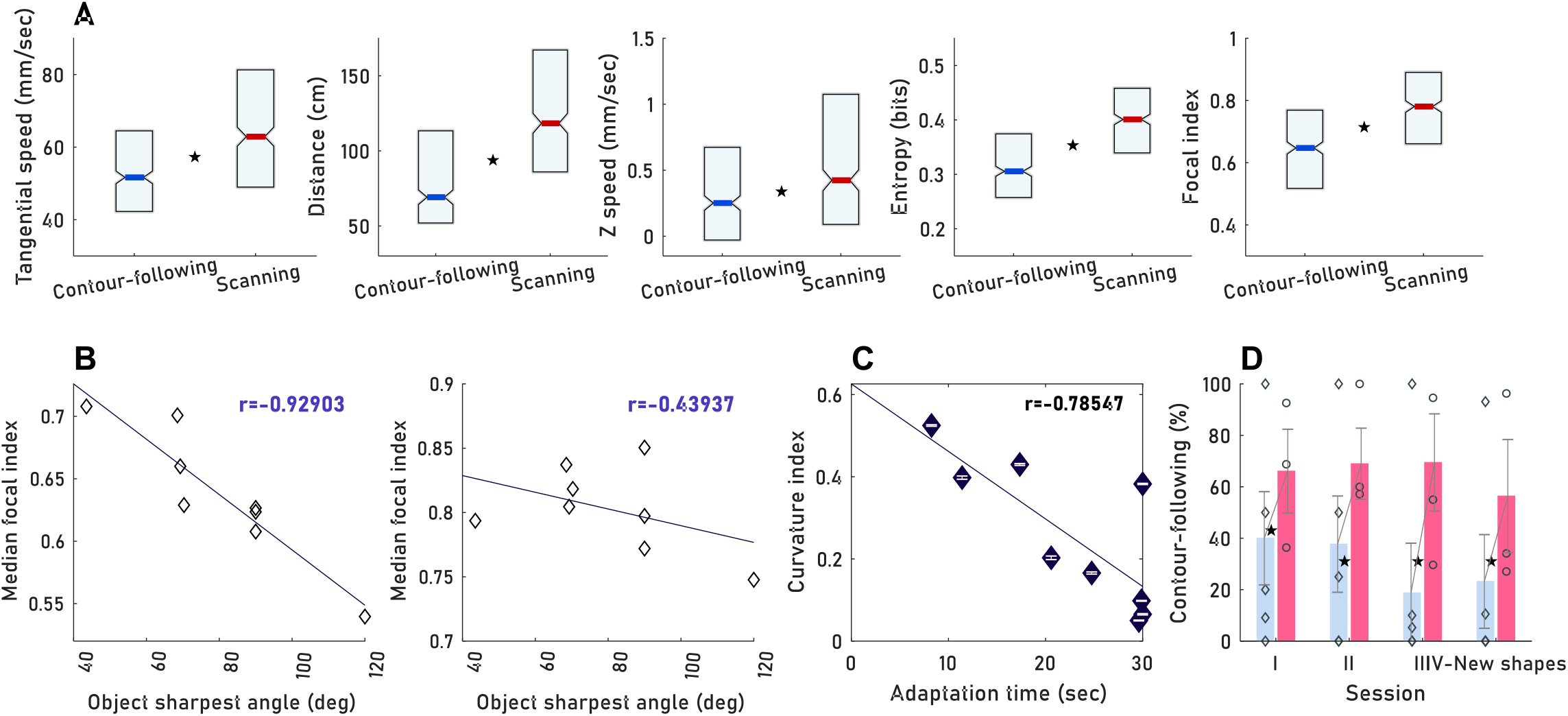
Results were replicated using human coder’s classification: **A** - *Sc* and *CF* classified by the human coders, differed in their mean kinematics and focal index (p<0.005), replicating the algorithm based differences (Fig. 2C, Fig. 3A). **B** - Objects median focal and objects sharpest angle are negatively correlated for human classified *CF* (left) but not for *Sc* trials (right). Replicating the correlation observed using the algorithm (Fig. 3C-D). **C** - Human classified *CF* trials curvature indices were highly correlated with participant’s adaptation times. (r=-0.78, p=0.012, adjusted alpha=0.01, 0.05/5), as in the case of the algorithm categorization (Fig. 4C). **D**-Prevalence of *CF* trials per session for groups I (light blue) and II (pink), group II used significantly more *CF* in all sessions (p<0.005) as in the case of the algorithm categorization (Fig. 5A).

**Figure 1-Sp5:**
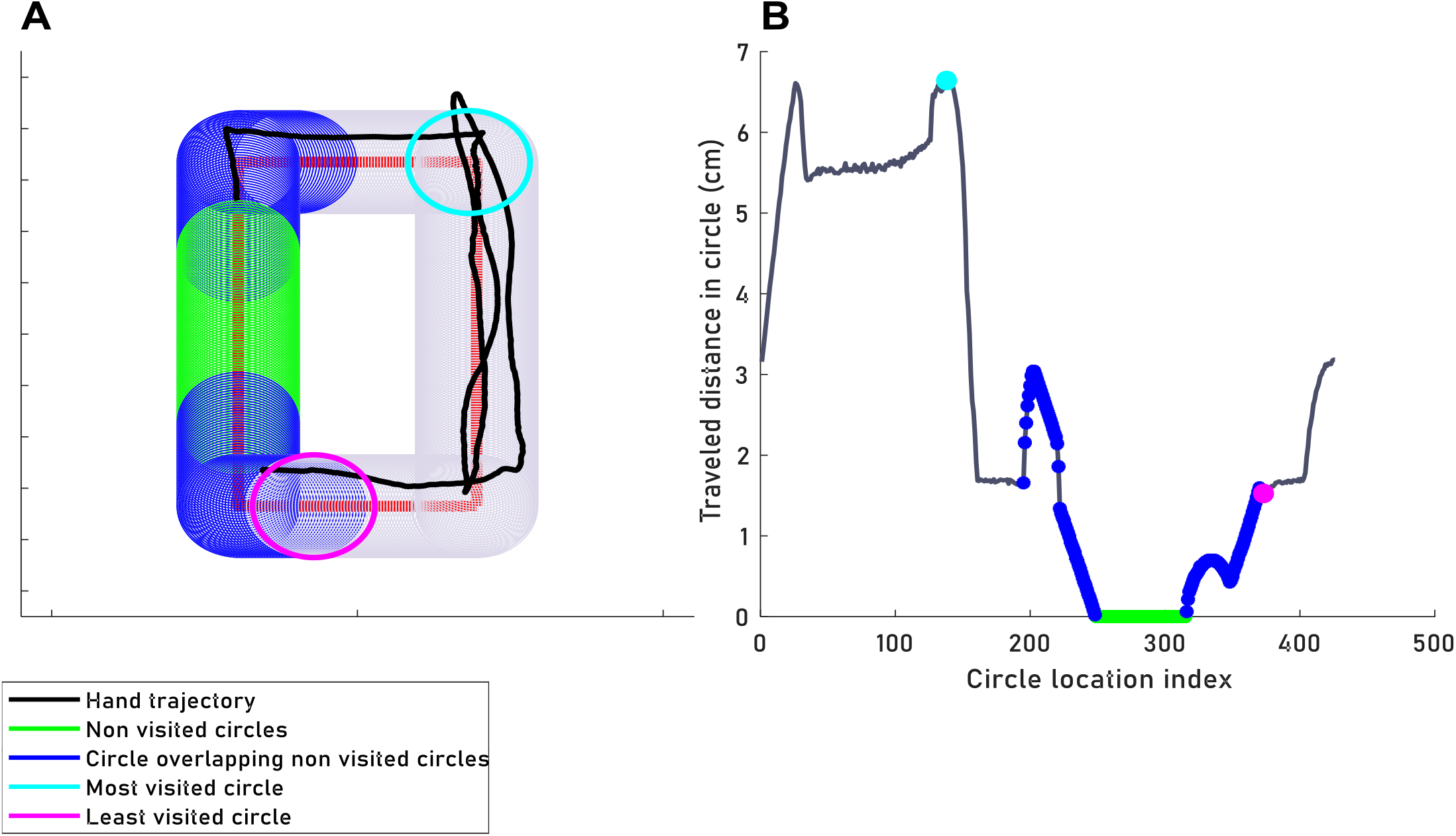
Removing zero circles or circles overlapping these zero circles: **A** - Overlapping circles with a radius of 10 mm (gray) were plotted on the shape’s outline (red). **B** - Calculated traveled distance (y-axis) in each of the circles (x-axis).The least visited circle (purple) was calculated after the exclusion of circles that were not visited at all (green), or circles that their area overlapped such circles (blue).

**Figure 1-Sp6:**
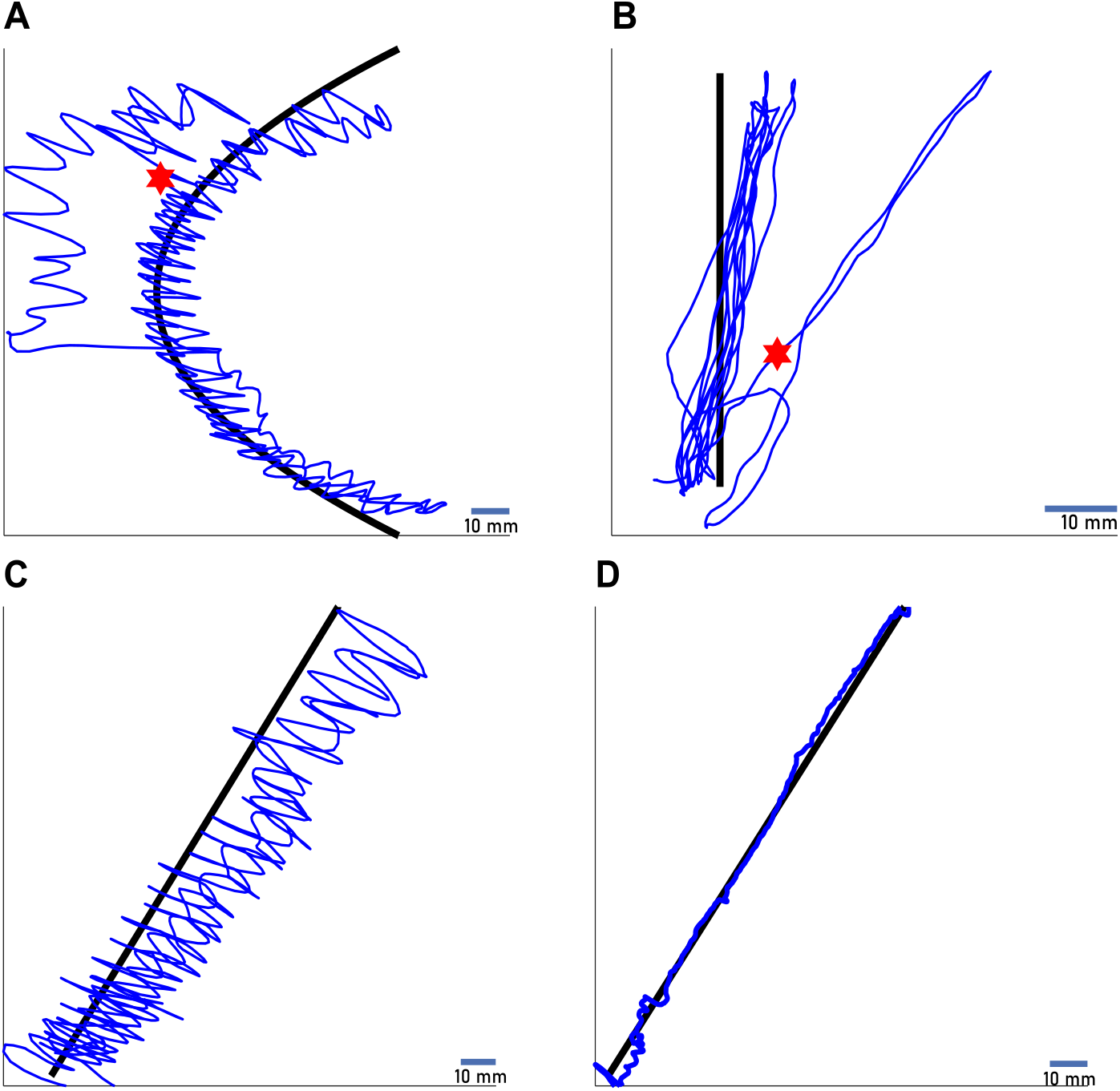
Examples of outline following trials: **A, B** - Hand trajectory of outline-following using *Oscillating* (A) or *Linear* (B) motion. The point of deviation from the outline is marked (red pentagram). The elapsed time from trial start to the deviation point is the trial τ_a_. **C, D** - Hand trajectory of outline following using *Oscillating* (C) or *Linear* (D) motion, in the case of no deviation from the outline. For these cases, Ta was assigned as the maximal trial duration (30 sec).

**Figure 1-Sp7:**
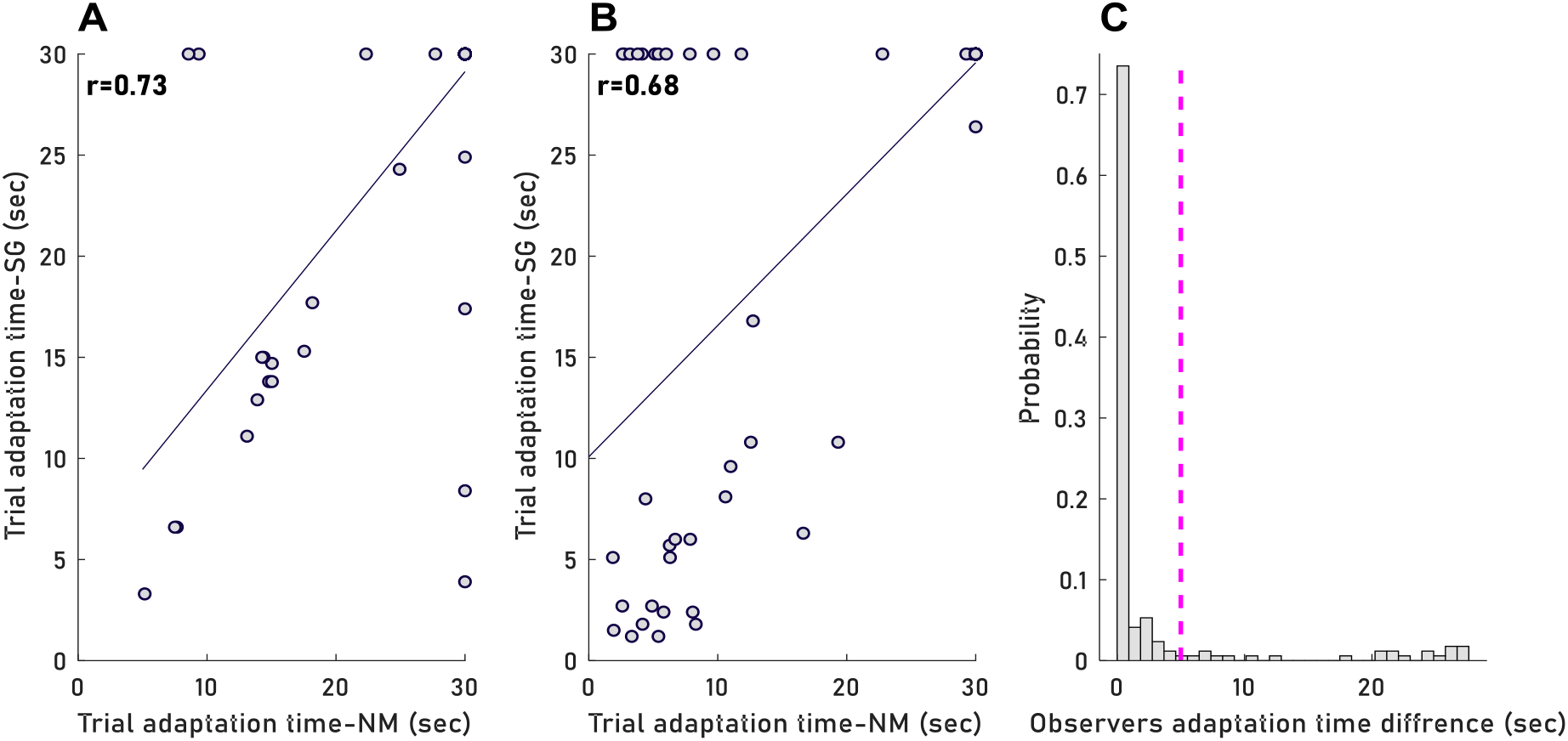
Observers correlations: **A, B** - Each data point represents the trial adaptation time marked by the two observers, for *Oscillating* trials (A) or *Linear* trials (B), p<0.005 for both conditions. **C** - Histogram of the differences between observer’s adaptation times in absolute value. Trials in which the difference was bigger than 5 sec (pink dashed line) were excluded.

**Figure 2-Sp8:**
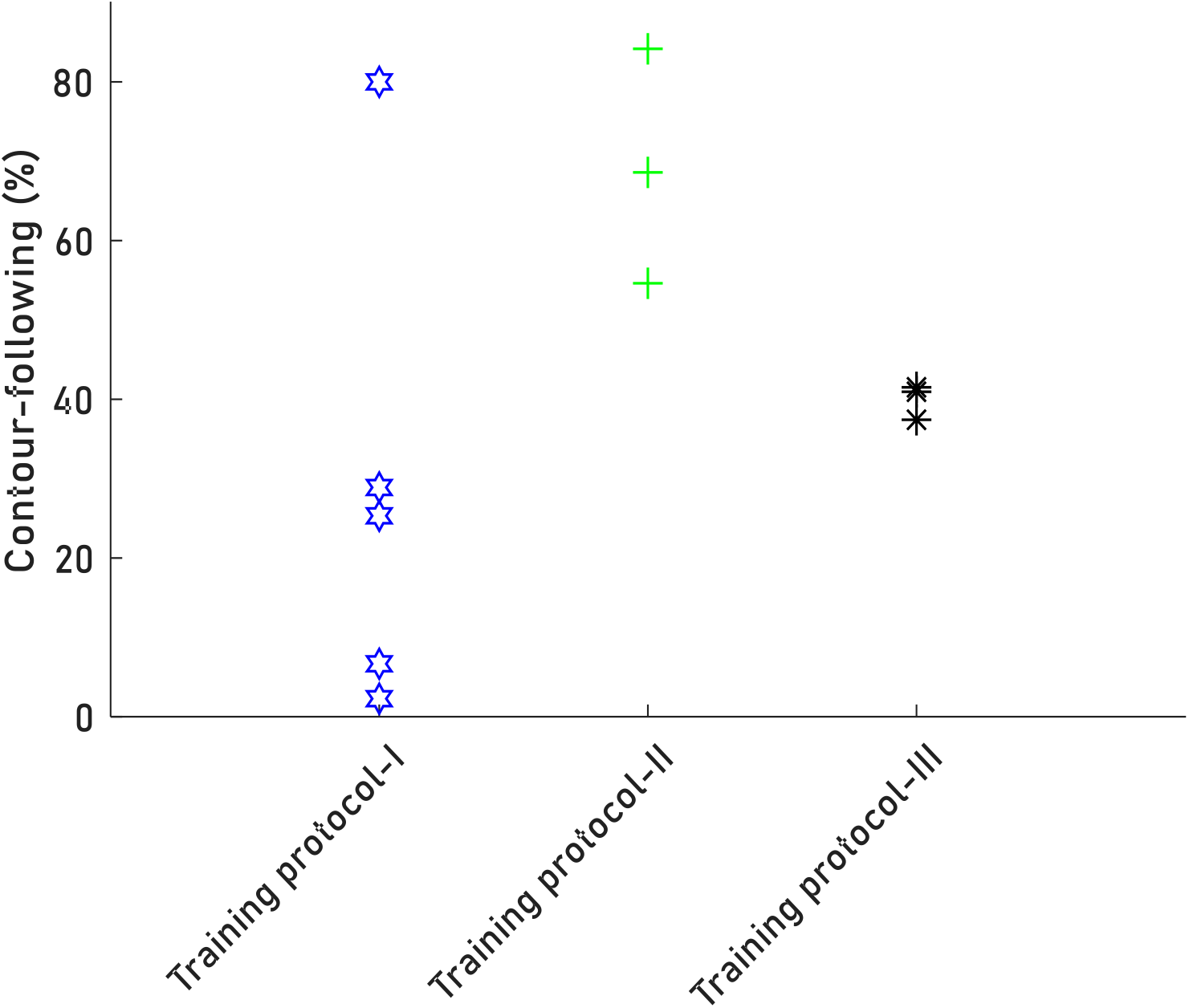
Precent of *CF* trials per protocol: Each data point represents the mean precent of *CF* trials per subject across all of her or his sessions. For protocol I (blue star), II (green cross) or III (black asterisk).

**Figure 2-Sp9:**
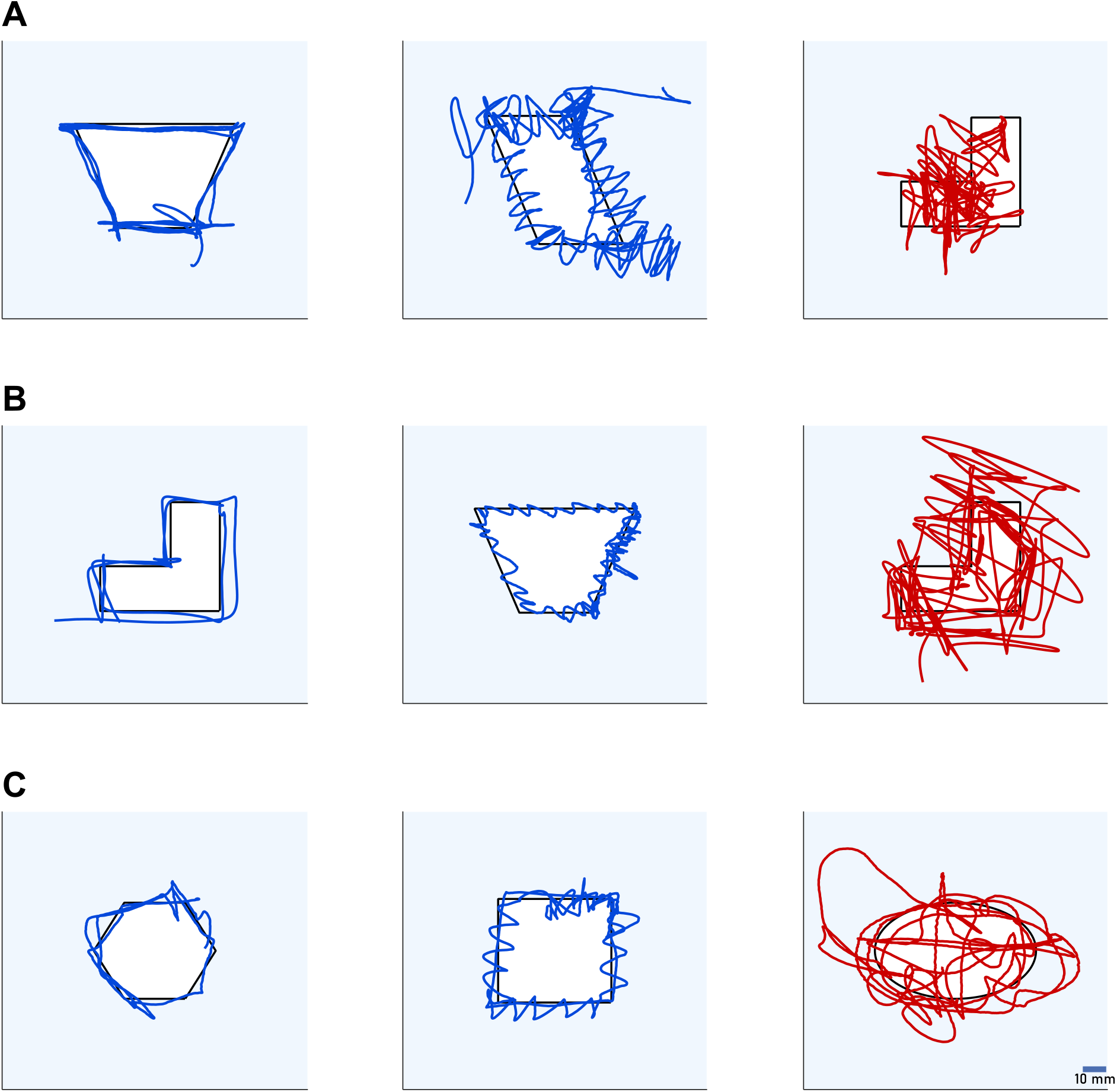
Example of *CF* and *Sc* trials for each training protocol: Examples of *Linear* and *Oscillating CF* trials (blue, left and middle columns, corresponding) and *Sc* trials (red, right column) for training protocol I (A), II (B) or III (C). Scale bar (C,right) stands for all example trials.

**Figure 4-Sp10:**
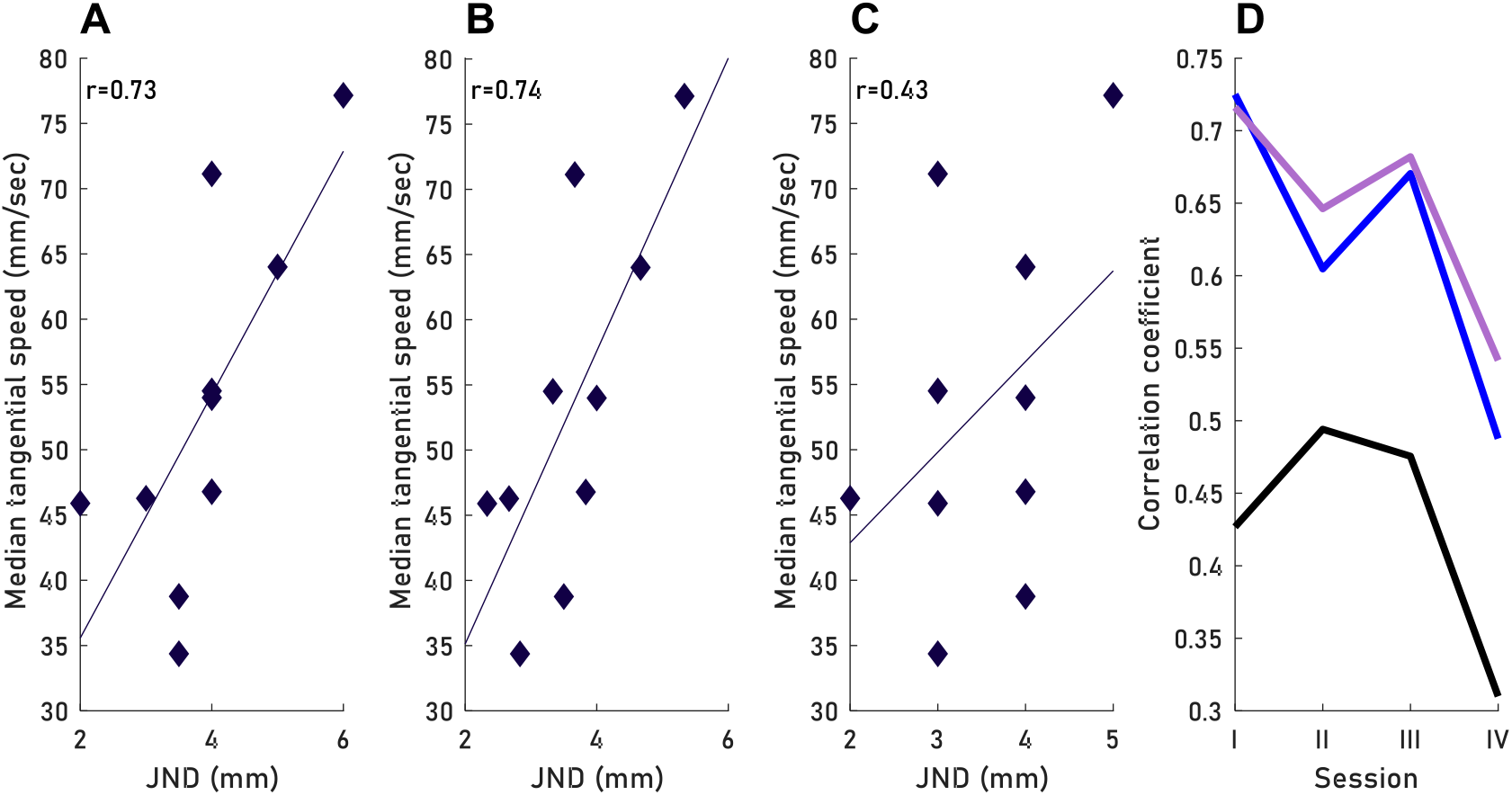
Dependency on spatial resolution in different fingers. **A,B,C** - Each data point represent one participant’s tangential speed across all trials of session one and its JND value at the index (A), mean JND value of the three fingers (B) or ring finger (C). **D** - Correlation coefficient between JND and tangential speed per session for either the index finger (blue), ring finger (black), or the mean JND value of all fingers (purple).

**Figure 4-Sp11:**
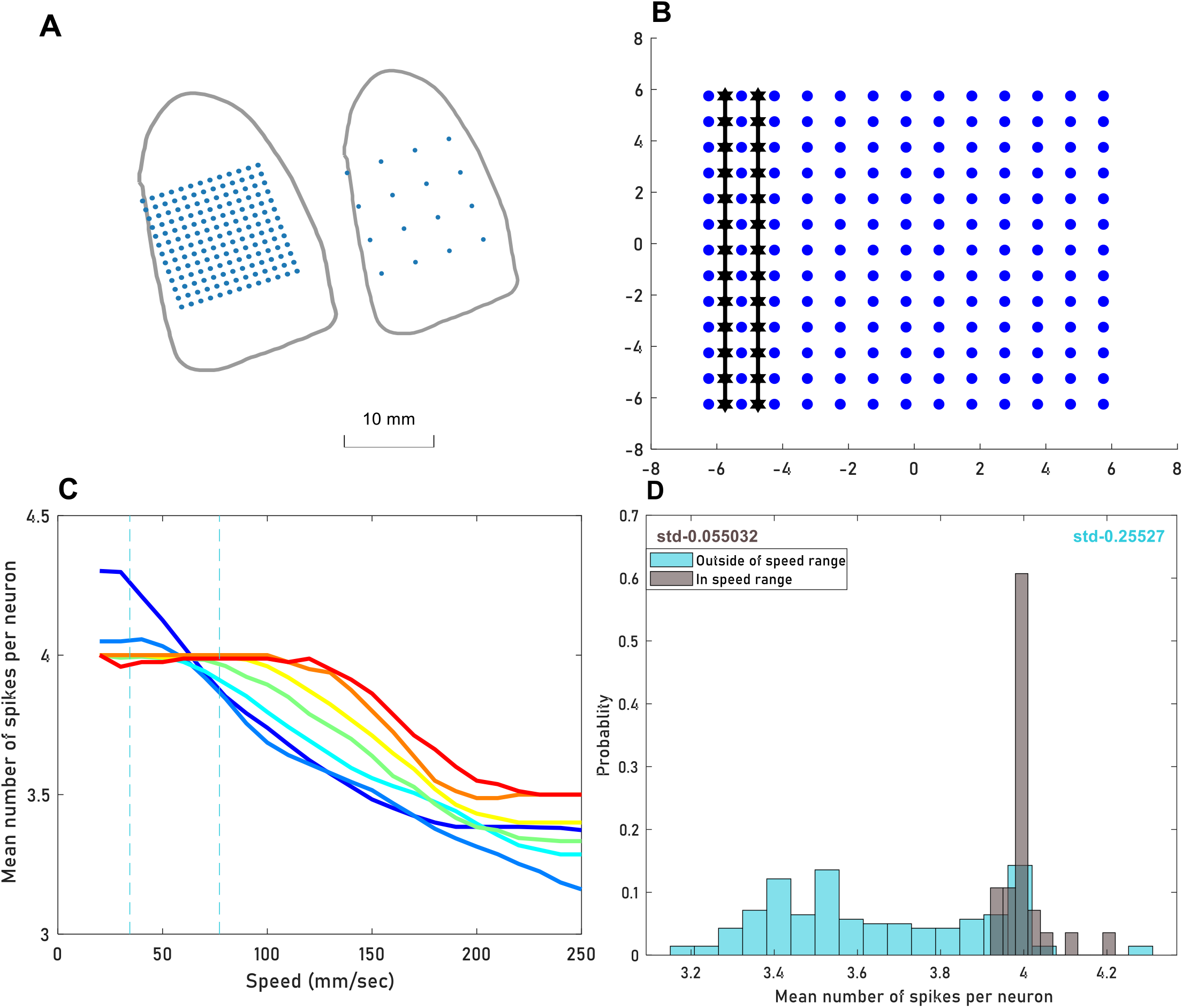
Spikes counts in relation to speed: **A** - An example of two finger grids, with either a 1 mm distance between units (left) or a 4 mm distance (right). **B** - Stimulus columns (black) were placed between the units’ columns (blue). In the presented grid the distance between units is 1 mm. **C** - The mean number of spikes per neuron plotted against speed. Color code represent the distance between units in the grid (blue for a small distance). The range of speed used by our participants is marked by the dashed gray lines. **D** - Histogram of spike counts per neuron, for speed values used by our participants (gray) or not (turquoise).

**Figure 4-Sp12:**
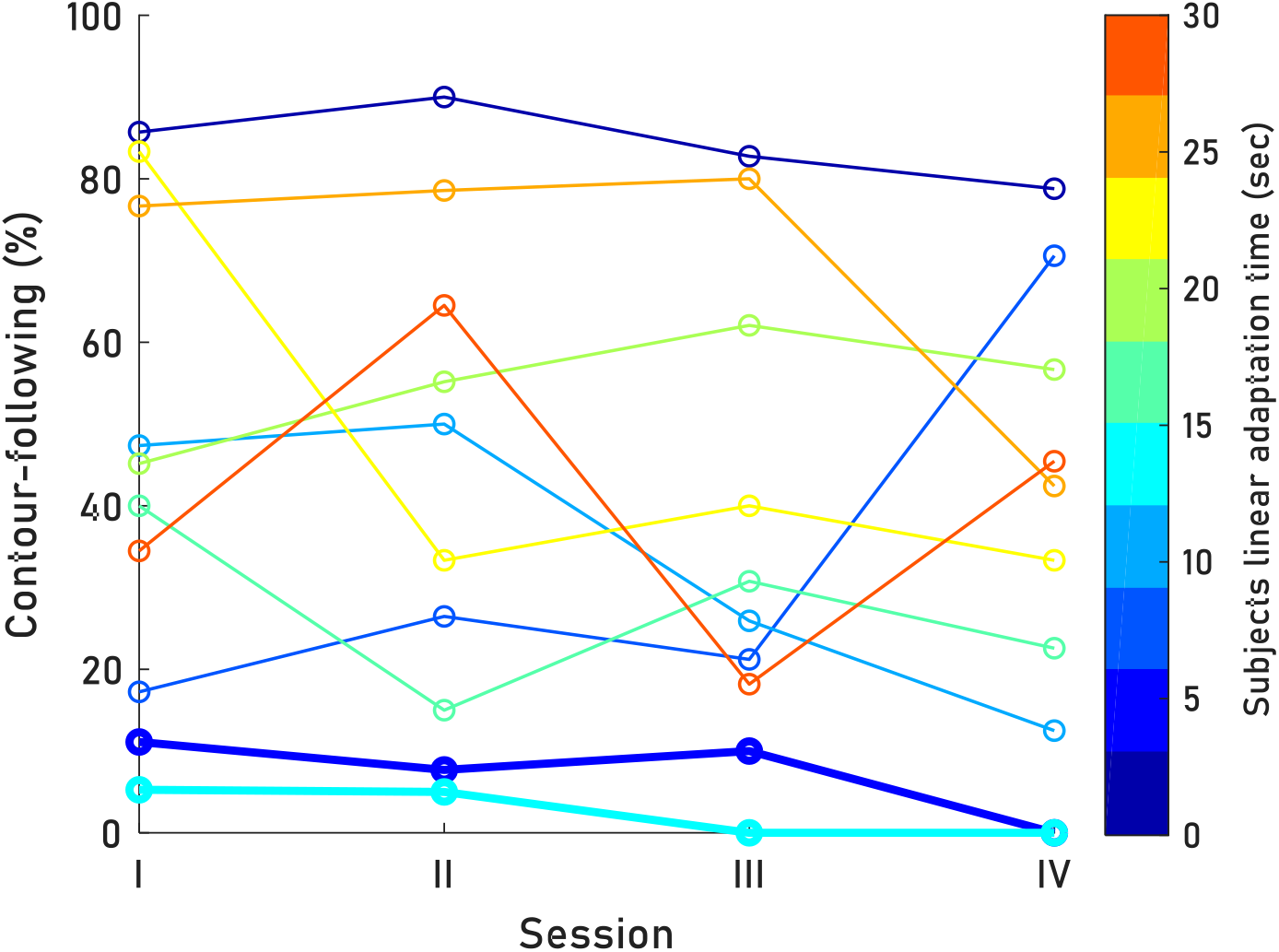
Changes in strategy choice between sessions: Each line represents the percent of *CF* trials per session of one subject. Color code stands for the participant *Linear* adaptation time. Two participants with short adaptation times (bold line) has stopped their use of *CF*.

**Table Sp1:**
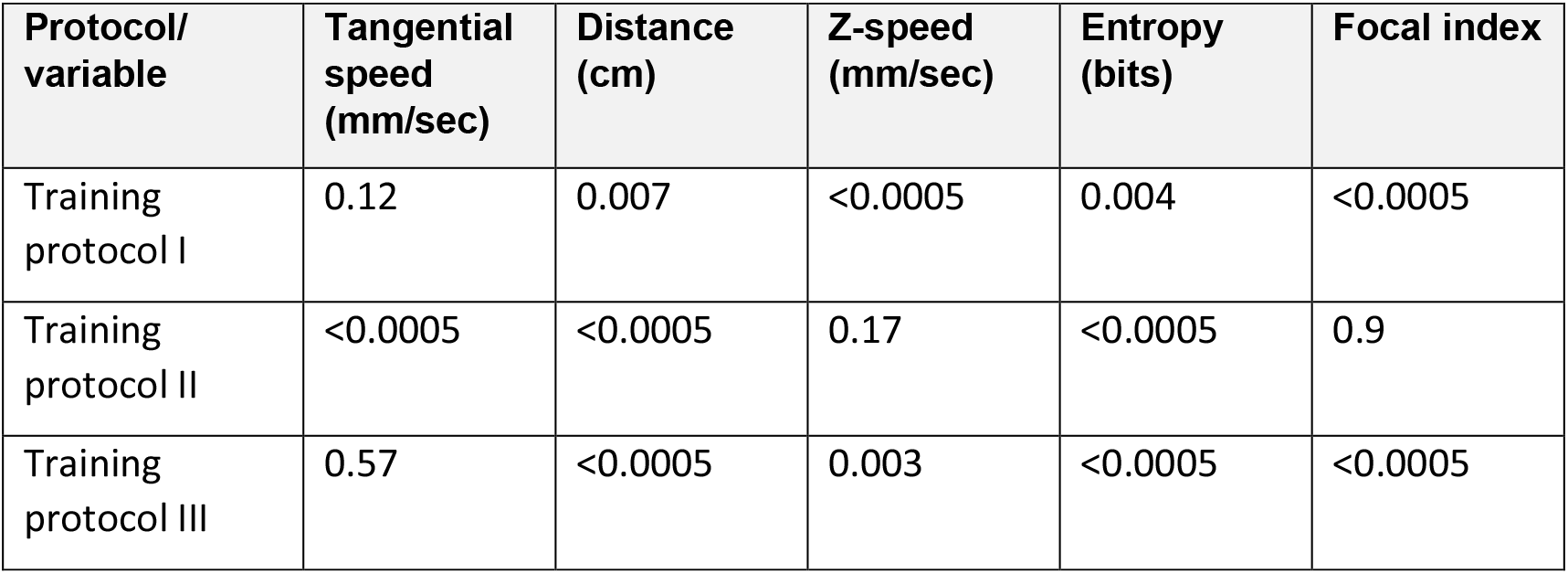
Kinematics differences for *CF* and Sc trials for each training protocol: p-value for each comparison between *Sc* and *CF* trials for each variable and training protocol. p-values are model, all significant p values (<0.05) were replicated using a bootstrap method.

| Protocol/<br>variable | Tangential<br>speed<br>(mm/sec) | Distance<br>(cm) | Z-speed<br>(mm/sec) | Entropy<br>(bits) | Focal index |
| --- | --- | --- | --- | --- | --- |
| Training<br>protocol I | 0.12 | 0.007 | <0.0005 | 0.004 | <0.0005 |
| Training<br>protocol II | <0.0005 | <0.0005 | 0.17 | <0.0005 | 0.9 |
| Training<br>protocol III | 0.57 | <0.0005 | 0.003 | <0.0005 | <0.0005 |

**Table Sp2:** Two-point discrimination JND per participant: JND values in a static two-point discrimination task in the pads of the middle, index and ring fingers, and mean values across these fingers

| Participant | Index finger<br>JND (mm) | Middle finger<br>JND (mm) | Ring finger<br>JND (mm) | Mean of three<br>fingers (mm) |
| --- | --- | --- | --- | --- |
| AF | 3 | 2 | 2 | 2.33 ±0.57 |
| EV | 3 | 3 | 4 | 3.33 ±0.57 |
| MF | 4 | 5 | 5 | 4.66 ±0.57 |
| AS | 3 | 4 | 4 | 3.66 ±0.57 |
| YB | 4 | 3.5 | 4 | 3.83 ±0.28 |
| SG | 5 | 5 | 6 | 5.33 ±0.57 |
| DR | 4 | 3 | 3.5 | 3.50 ±0.50 |
| HA | 4 | 4 | 4 | 4 ±0 |
| MG | 2 | 3 | 3 | 2.66 ± 0.57 |
| ZA | 3 | 2 | 3.5 | 2.83 ± 0.76 |

## Notes

**Conflict of interest statement:** The authors declare no competing financial interests.

### Competing Interest Statement

The authors have declared no competing interest.

